# Evolution in alternating environments with tunable inter-landscape correlations

**DOI:** 10.1101/803619

**Authors:** Jeff Maltas, Douglas M. McNally, Kevin B. Wood

## Abstract

Natural populations are often exposed to temporally varying environments. Evolutionary dynamics in varying environments have been extensively studied, though understanding the effects of varying selection pressures remains challenging. Here we investigate how cycling between a pair of statistically related fitness landscapes affects the evolved fitness of an asexually reproducing population. We construct pairs of fitness landscapes that share global fitness features but are correlated with one another in a tunable way, resulting in landscape pairs with specific correlations. We find that switching between these landscape pairs, depending on the ruggedness of the landscape and the inter-landscape correlation, can either increase or decrease steady-state fitness relative to evolution in single environments. In addition, we show that switching between rugged landscapes often selects for increased fitness in both landscapes, even in situations where the landscapes themselves are anti-correlated. We demonstrate that positively correlated landscapes often possess a shared maximum in both landscapes that allows the population to step through sub-optimal local fitness maxima that often trap single landscape evolution trajectories. Finally, we demonstrate that switching between anti-correlated paired landscapes leads to ergodic-like dynamics where each genotype is populated with nonzero probability, dramatically lowering the steady-state fitness in comparison to single landscape evolution.

## I. INTRODUCTION

Natural populations experience tremendous environmental diversity, and understanding how this spatiotemporal diversity influences evolutionary dynamics is a long-standing challenge. A great deal of work, both theoretical and experimental, has shown that spatial (Agarwala and Fisher 2019; Constable and McKane 2014a,b; Farhang-Sardroodi et al. 2017; Habets et al. 2006; Hermsen and Hwa 2010; Korona et al. 1994; Lin et al. 2015; Waddell et al. 2010; Whitlock and Gomulkiewicz 2005) and temporal (Acar et al. 2008; Canino-Koning et al. 2019; Cook and Hartl 1974; Cooper and Lenski 2010; Cvijović et al. 2015; Gaál et al. 2010; Gillespie and Guess 1978; Gupta et al. 2011; Hartl and Cook 1974; Kashtan et al. 2007; Kussell and Leibler 2005; Lewontin and Cohen 1969; Mustonen and Lässig 2008; Mustonen and Lässig 2009; Patra and Klumpp 2015; Shahrezaei et al. 2008; Skanata and Kussell 2016; Steinberg and Ostermeier 2016; Tan and Gore 2012; Tan et al. 2011; de Vos et al. 2015) heterogeneity play an important role in adaptation of asexual communities. For example, temporal or spatial fluctuations may lead to increased fixation probability and adaptation rates (Cvijović et al. 2015; Farhang-Sardroodi et al. 2017; Hermsen and Hwa 2010; Kashtan et al. 2007; Lewontin and Cohen 1969; Mustonen and Läassig 2008; Whitlock and Gomulkiewicz 2005), a phenomenon that is also exploited in genetic programming algorithms (ONeill et al. 2010). In addition, environments that change in systematic ways may promote facilitated variation (Gerhart and Kirschner 2007; Parter et al. 2008), allowing organisms to preferentially harness the beneficial effects of random genetic changes and rapidly adapt to future perturbations. And when phenotypes themselves fluctuate over time, the frequency of inter-phenotype switching can evolve to match the timescale of environmental fluctuations (Acar et al. 2008; Gupta et al. 2011; Kussell and Leibler 2005; Shahrezaei et al. 2008).

It is increasingly clear that these evolutionary dynamics have practical consequences for human health. The rise of drug resistance, which threatens the efficacy of treatments for bacterial infections, cancer, and viruses, is driven–at least in part–by evolutionary adaption occurring in complex, heterogeneous environments. Spatial heterogeneity in drug concentration has been shown to accelerate the evolution of resistance (Baym et al. 2016; Fu et al. 2015; Greulich et al. 2012; Hermsen et al. 2012; Moreno-Gamez et al. 2015; Zhang et al. 2011), though adaptation may also be slowed when fitness landscapes (Greulich et al. 2012) or drug profiles (De Jong and Wood 2018) are judiciously tuned. Similarly, temporal variations in drug exposure–for example, drug cycling–can slow resistance under some conditions, though hospital-level strategies such as mixing may be more effective at generating the requisite environmental heterogeneity (Bergstrom et al. 2004; Brown and Nathwani 2005). Recent studies have also shown the potential of new control strategies that harness so-called *collateral effects* (Barbosa et al. 2018; Dhawan et al. 2017; de Evgrafov et al. 2015; Fuentes-Hernandez et al. 2015; Imamovic et al. 2018; Imamovic and Sommer 2013; Kim et al. 2014; Lazar et al. 2018, 2014, 2013; Maltas et al. 2019; Maltas and Wood 2019; Munck et al. 2014; Nichol et al. 2019; Roemhild et al. 2015, 2018; Yoshida et al. 2017), which occur when resistance to a target drug is accompanied by an increase or decrease in resistance to an unseen stressor. In essence, these strategies force populations to simultaneously adapt to incompatible evolutionary tasks (Hart et al. 2015; Shoval et al. 2012).

Evolutionary adaptation is often modeled as a biased random walk on a high-dimensional landscape that links each specific genotype with a particular fitness (Gillespie 1983a,b, 1984). In the simplest scenario, these landscapes represent evolution in the strong selection weak mutation (SSWM) limit, where isogenic populations evolve step-wise as the current genotype is replaced by that of a fitter descendant. While these idealized models are strictly valid only under certain conditions–for example, SSWM typically holds when mutation rate and effective population size are small–simple models have contributed significantly to our understanding of evolution (Cook and Hartl 1974; Desai and Fisher 2007; Desai et al. 2007; Gerrish and Lenski 1998; Gillespie 1983a, 1984; Hartl and Cook 1974). In the context of fitness landscape models, control strategies that exploit collateral effects force the population to adapt to sequences of distinct, but statistically related, landscapes. For example, alternating between two drugs that induce mutual collateral sensitivity (adaptation to drug A leads to sensitivity to drug B, and vice versa) corresponds to landscapes with anti-correlated fitness peaks. When environments change in systematic ways–for example, by forcing the population to adapt to modular tasks comprised of related sub-goals–adaptation may select for generalists, genotypes that are fit in a wide range of environments at the cost of suboptimal specialization for any particular task (Parter et al. 2008; Sachdeva et al. 2020; Wang and Dai 2019). Relatively recent theoretical work also shows that conditional effects of evolutionary history can be captured by slowly changing landscapes–*seascapes*–which allow for the incorporation of time-dependent correlations (Agarwala and Fisher 2019; Mustonen and Lässig 2009). In general, however, understanding evolution in correlated landscapes–and in particular, how the choice of that correlation impacts fitness adaptation–remains challenging.

In this work, we investigate evolutionary dynamics of asexual populations in rapidly alternating environments described by pairs of (potentially rugged) fitness landscapes with tunable inter-landscape correlations (Fig 1). This problem is loosely inspired by adaptation of microbial communities to 2-drug cycles in which each drug induces collateral resistance or sensitivity to the other, though the scenario in question may arise in many different contexts, including evolution in antibodies (Burton et al. 2012) and viruses (Rhee et al. 2010). Our goal is to understand how the interplay between intra-landscape disorder (ruggedness) and inter-landscape fitness correlations impact fitness. By formulating the evolutionary dynamics as a simple Markov chain (Durrett and Durrett 1999; Nichol et al. 2015), we are able to efficiently calculate time-dependent genotype distributions and investigate adaptation to ensembles of landscape pairs with various levels of epistasis and fitness correlations–results that would be more difficult to achieve from stochastic simulations alone. We find that rapid switching can either increase or decrease the steady state fitness of the population, depending on both the correlation between landscapes and level of intra-landscape ruggedness (i.e. epistasis). On short timescales, mean fitness is generally highest in static landscapes, but rapid switching between correlated environments can produce fitness gains for sufficiently rugged landscapes on longer timescales. Surprisingly, longer periods of rapid switching can also produce a genotype distribution whose fitness is, on average, larger than that of the ancestor population in both environments, even when the landscapes themselves are anticorrelated. To intuitively understand these results, we visualized genotype distributions and inter-genotype transitions as network diagrams, revealing that rapid switching in highly correlated environments frequently shepherds the population to genotypes that are locally optimal in both landscapes and, in doing so, fosters escape from the locally optimal but globally suboptimal fitness peaks that limit adaptation in static environments. The dynamics arise, in part, from the fact that rugged landscape pairs are increasingly likely to exhibit shared maxima as they become more positively correlated, and in turn, for landscapes with positive correlations, the mean fitness of these shared peaks is higher than that of nonshared peaks. By contrast, evolution in anti-correlated landscape pairs sample large regions of genotype space, exhibiting ergodic-like steady-state behavior that results in decreased average fitness.

**FIG. 1:**
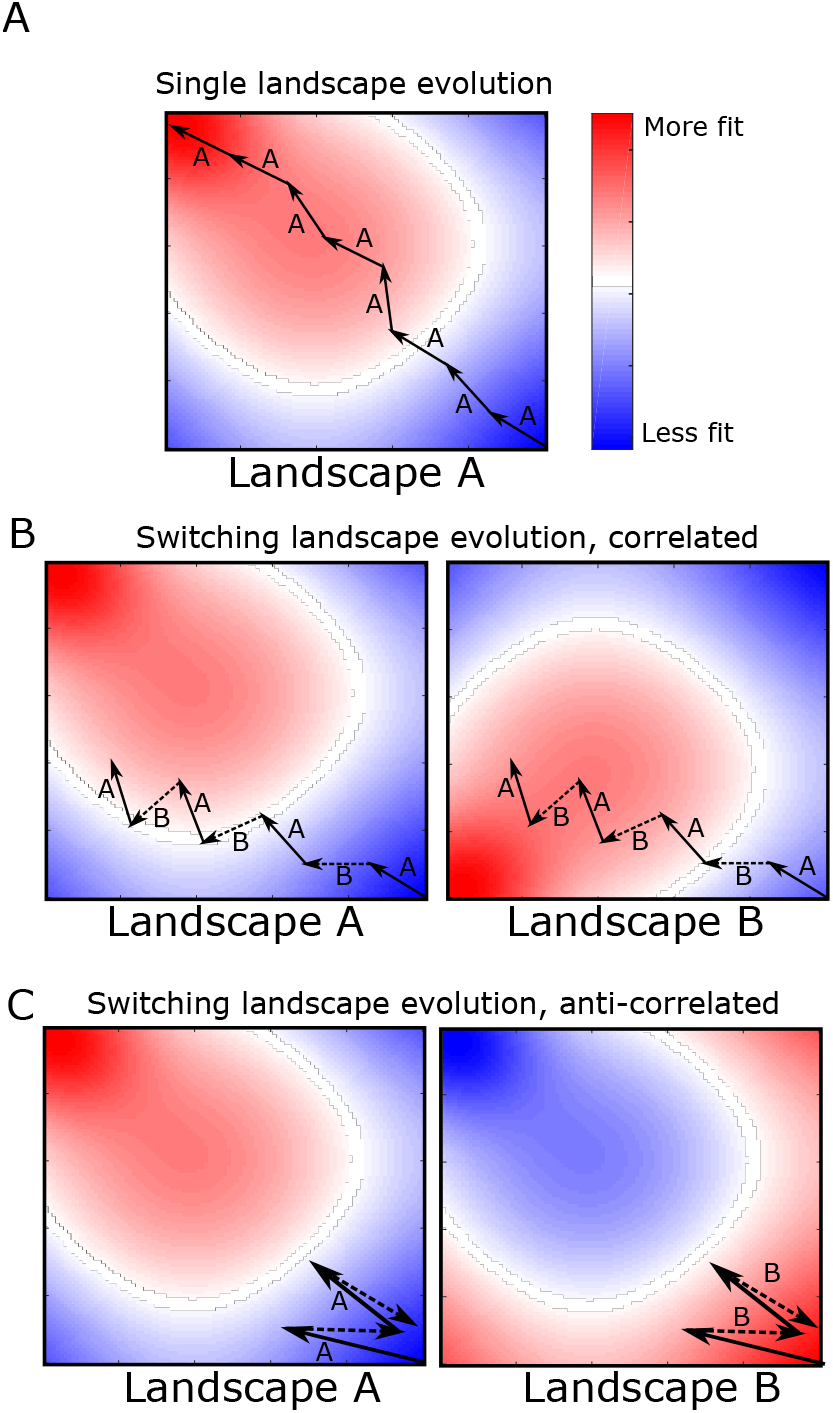
Adaptation to alternating landscapes may depend on inter-landscape correlations. A. Schematic fitness landscape, with fitness varying from less fit (blue) to more fit (red) over the two dimensional genotype space. Starting from a single genotype (lower right hand corner), adaptation follows a biased random walk (arrows) toward local fitness maxima (in this case, in the upper left side of the landscape). B and C. Fitness landscapes A and B are positively (B) or negatively (C) correlated and do not share a global fitness maximum. Adaptation under rapid alternation of landscapes A and B leads to an altered evolutionary trajectory (represented as arrows, with solid arrows indicating steps in A and dashed arrows steps in B). In this example, the final fitness achieved in both correlated (panel B) and anti-correlated (panel C) landscapes is lower than that of static landscape evolution (panel A). Adaptation to anti-correlated landscapes leads to a particularly significant decrease in final fitness, as each step in B effectively reverses the progress made the previous step in A.

## II. RESULTS

### A. Markov chain model of evolution in alternating landscape pairs with tunable correlations

We consider evolution of an asexual haploid genome with *N* mutational sites. Each mutational site can have one of two alleles (labeled 0 or 1), and a single genotype can therefore be represented by one of the 2^*N*^ possible binary sequences of length N. The fitness of each genotype depends on the specific environment in which evolution takes place. We consider two different environments (“A” and “B”), and in each environment, every genotype is assigned a fixed fitness value, which defines the corresponding fitness landscapes (landscape A and landscape B) in each environment. Each fitness landscape is therefore defined on an *N*-dimensional hypercubic graph, with the nodes corresponding to specific genotypes.

To construct the landscape for a given environment, we use a many-peaked “rough Mt. Fuji” landscape (Aita and Husimi 1998; Neidhart et al. 2014; Tan and Gore 2012). Specifically, we assume that the fitness of the ancestor genotype (0,0,0…0) is zero and that the fitness *f_i_* associated with a single mutation at mutational site *i* is drawn from a uniform distribution on the interval [-1,1]. Single mutations can therefore lead to increases (*f_i_* > 0) or decreases (*f_i_* < 0) in fitness. To fully specify the base landscape (i.e. the smooth landscape in the absence of epistasis), we then assume fitness associated with multiple mutations is additive. Finally, landscape ruggedness is incorporated by adding to the fitness of each genotype *j* a fixed, random variable *ξ_j_* drawn from a zero-mean normal distribution with variance *σ*^2^. The variable *σ*–the amplitude of the noise–determines the level of ruggedness of the landscape, which simulates epistasis (Anderson et al. 2015; Phillips 2008; Ritchie et al. 2001; da Silva et al. 2010; Tsai et al. 2007; Weinreich et al. 2006; Xu et al. 2005). In what follows, we focus on landscapes of size *N* = 7 (128 total genotypes) for computational convenience and limit ourselves primarily to *σ* = 0 (smooth landscapes) or *σ*=1 (rugged landscapes).

Our goal is to investigate evolution in rapidly changing environments that correspond to landscape pairs with correlated fitness peaks. To do so, we generate for each landscape A a “paired” landscape B with similar statistical properties (identical fitness mean and variance) but fitness peaks that are, on average, correlated with those of landscape A in a tunable way. To do so, we represent each landscape A as a vector 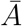 of length 2^*N*^ and use simple matrix algebra to generate a random vector 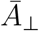 orthogonal to 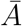; by construction, then, this vector corresponds to a landscape whose fitness values are, on average, uncorrelated with those of landscape A. It is then straightforward to generate a vector 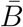, a linear combination of 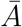 and 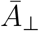, such that the fitness values of landscapes A and B are correlated to a tunable degree −1 ≤ *ρ* ≤ 1, where *ρ* is the Pearson correlation coefficient between the two vectors 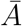 and 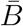 (see Methods).

With the landscapes specified, we then model adaptation in the well-characterized Strong Selection Weak Mutation (SSWM) limit (Gillespie 1983a,b, 1984), which can be formally described by a Markov chain (Durrett and Durrett 1999; Nichol et al. 2015). During each time step, the population transitions with uniform probability to one of the neighboring genotypes with a higher fitness in the current environment. We compare adaptation on a single landscape (single landscape evolution, SLE) with adaptation to rapid alternation of the two correlated landscapes A and B, which we refer to as paired landscape evolution (PLE). We focus here on the limit of rapid environmental switching, where the fitness landscape changes (A-B-A-B…) at each time step. This corresponds loosely to the rapid environmental switching seen in many laboratory experiments (Burch and Chao 1999; Crill et al. 2000; Kim et al. 2014; Lenski 1988).

We are primarily interested in comparing the (average) steady-state fitness of populations undergoing SLE to that of populations undergoing PLE. The average fitness, 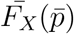, in environment *X* can be calculated at any time step *t* using 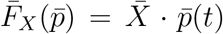, where 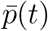 is the vector whose *i*^th^ component is the probability to be in genotype *i* at time t and 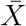 is the landscape vector for environment *X*. Because the process can be described by a Markov chain, the vector 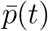 is given by 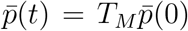, where the matrix *T_M_* describes the sequence of environments over time (e.g. 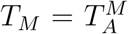 for *M* steps in environment *A*, or *T_M_* = (*T_B_T_A_*)^*M*/2^ for *M* consecutive A-B cycles, with *T_A_* and *T_B_* the transition matrices corresponding to single steps in environment A and B, respectively). In what follows, we focus primarily on the mean fitness difference between the SLE and PLE adaptation, which is given by 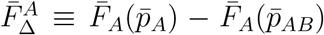, where 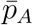 is the steady state genotype distribution following adaptation to environment A, and 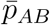 is the steady state genotype distribution following adaptation to alternating A-B environments. Note that we define this fitness difference, 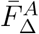, with respect to landscape 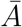 (noted by superscript), which allows us to compare adaptation in environment A with adaptation in the alternating A-B environments. In the drug cycling analogy, we are measuring the average fitness in the drug A environment–essentially a measure of resistance to that drug. In all calculations, we consider an ensemble of 1000 landscapes pairs–with each pair sharing the same mean and variance in fitness and the same inter-landscape correlations–and we average the results over this ensemble.

## B. Adaptation in rugged landscapes frequently ends in local, sub-optimal fitness maxima

While adaptation to static, rugged landscapes is well-understood, we first briefly discuss the effects of landscape ruggedness in the context of the current model. In static landscapes, steady state is reached when the genotype corresponds to a local fitness maximum. In the case of smooth, purely additive landscapes (*σ* = 0), there is a single fitness peak that corresponds to the global maximum, which we call gMax. However, as the landscape becomes more rugged (*σ* > 0), the average number of local maxima increases. For small *σ* ≪ 1, the average number of local maxima is *N_max_* ≈ 1 + 1/2*N*(*N* + 1)*σ*^2^, while for large *σ* it approaches the theoretical maximum of 2^*N*^/(*N* + 1) (Fig 2A); in the SI, we provide a semi-analytical approximation for intermediate values of *σ*. In turn, the fraction of adaptation trajectories that reach the global maximum decreases, reflecting the propensity of rugged landscapes to trap evolution in globally sub-optimal genotypes. To visualize these results, we represented the steady state genotype distributions and inter-genotype transitions as a network diagram (Fig 2B), with each node (circle) representing a genotype. The shading of each circle represents the relative fitness of that genotype (ranging from less fit, white, to more fit, black) and the size of the circle indicates occupation probability in the steady state. Arrows connecting different genotypes indicate nonzero transition probabilities, with the thickness of the arrow corresponding to its magnitude. We show only those transitions that can occur when adaptation starts in the ancestor genotype (top of diagram). In the case of evolution on a smooth landscape (*σ* = 0, Fig 2B, left panel), all trajectories lead to the single global maximum (indicated by red “+”). However, in the rugged landscape (*σ* = 1, Fig 2B, right panel), there is a nonzero probability of settling in each of three local maxima, and the population frequently ends in a non-optimal genotype. Increasing ruggedness would therefore be expected to lower the average fitness achieved in an ensemble of landscapes.

**FIG. 2:**
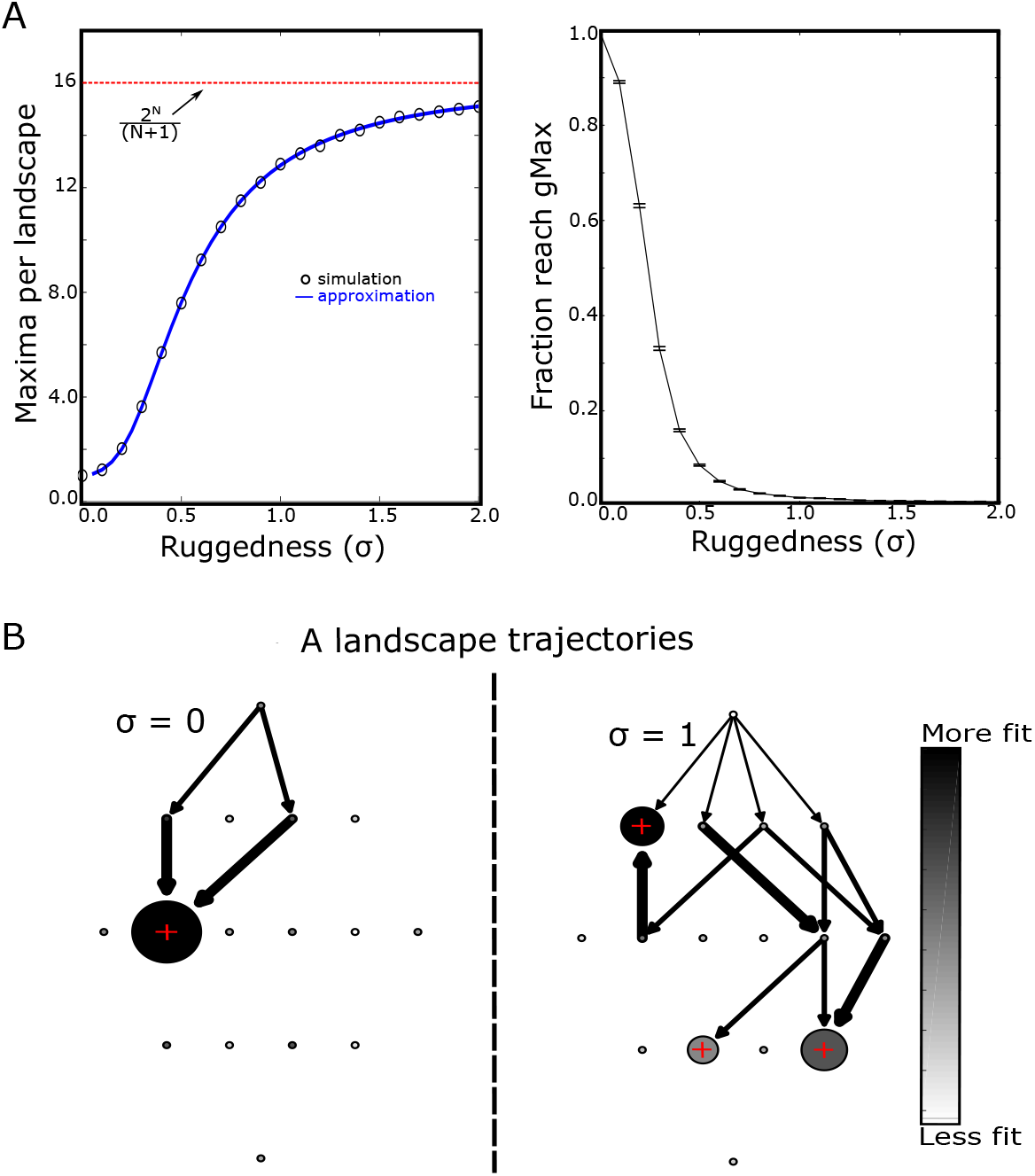
Rugged landscapes trap populations in non-optimal fitness maxima. A. Left panel: average number of local fitness maxima per landscape as a function of increasing ruggedness (epistasis, *σ*). Circles are estimates from simulations, solid curve is semi-analytical approximation (see SI), and dotted red line is the theoretical maximum (2^*N*^/(*N* + 1) = 16). Right panel: fraction of adapted populations that reach the global fitness maximum value as a function of ruggedness. Error bars are ± standard error of the mean in the ensemble of landscapes. B. Sample adaptive trajectories for small landscapes (*N* = 4) and *σ* = 0 (left) or *σ* = 1 (right). Each circle represents a genotype, with the ancestral genotype at the top. The shading of the circle represents the relative fitness of that genotype (ranging from less fit, white, to more fit, black) and the size of the circle indicates occupation probability in the steady state. Red + symbols mark genotypes corresponding to local fitness maxima. Arrows represent transitions between genotypes that occur with nonzero probability given that adaptation begins in the ancestral genotype.. The width of the arrow represents the magnitude of the transition probability.

## C. Switching between positively correlated landscapes can produce higher average fitness than adaptation to a static environment

Next, we set out to compare adaptation to landscape A with adaptation to alternating landscapes (A, B) with a tunable level of correlation, *ρ*, in the absence of epistasis (*σ* = 0, Fig 3A, blue). On these smooth landscapes, the fitness is single-peaked (Tan and Gore 2012), and in the absence of switching, the population always reaches this global maximum. In alternating environments, adaptation approaches the same average fitness as in static environments (i.e. 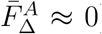)–implying that it finds the global fitness maximum–for all but the most negatively correlated landscapes (*ρ* < −0.85), where switching leads to steep decreases in fitness. By contrast, when landscapes are rugged (*σ* = 1), we find a range of correlations for which switching (PLE) increases the mean fitness (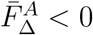, Fig 3A, orange). Furthermore, as ruggedness increases, the range of correlations leading to increased fitness grows (Fig 3B).

**FIG. 3:**
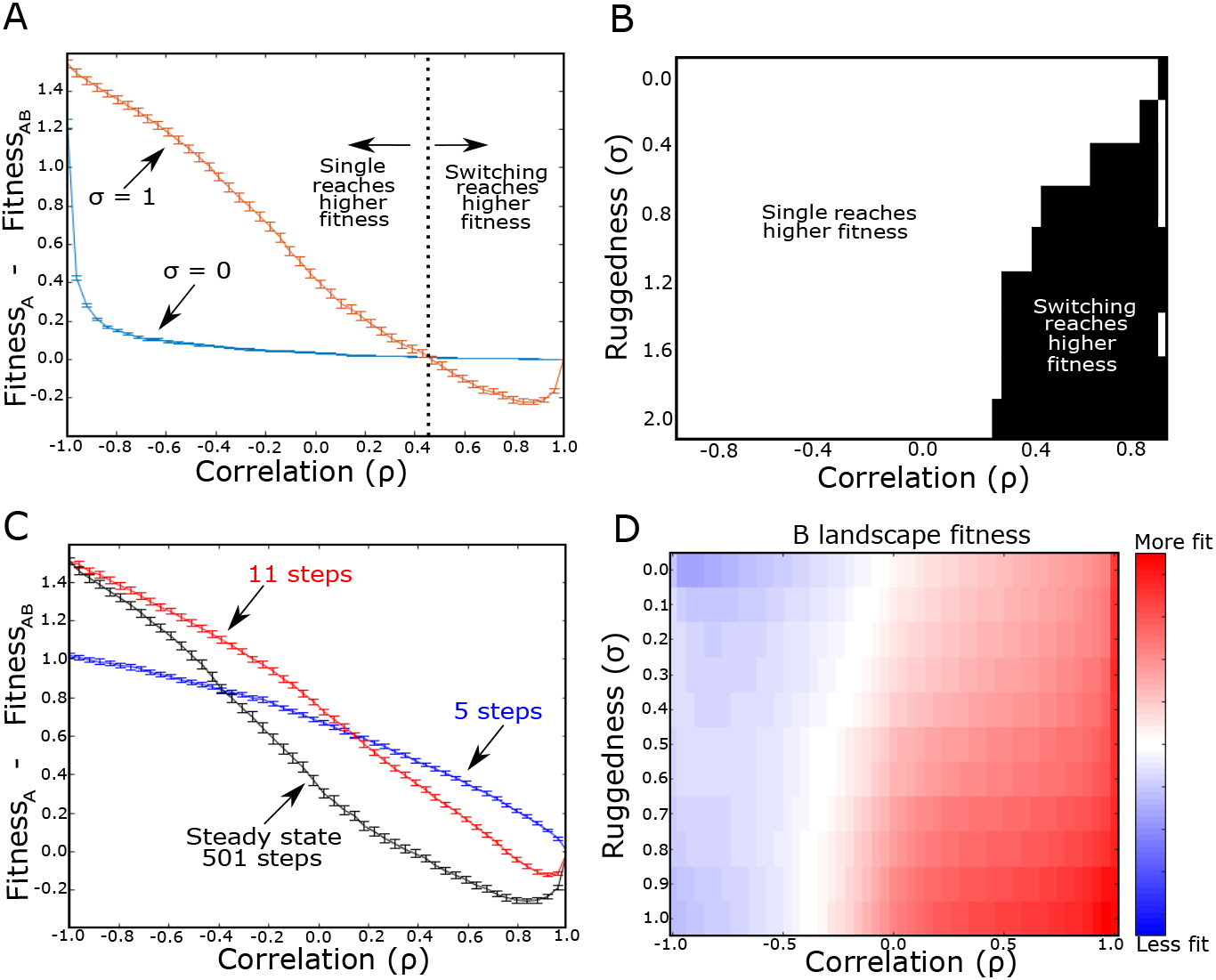
Modulated fitness in alternating landscapes depends on intra-landscape ruggedness and inter-landscape correlations. A. Difference in average fitness (at steady state) between populations adapted to a single static landscape (A) or rapidly alternating landscape pairs (A-B) as a function of correlation between landscapes A and B. Average fitness is defined as the mean fitness of the steady state genotypte distribution (which arises following adaptation to either static or switching protocols) measured in landscape A. Blue curve: *σ* = 0 (no epistasis; smooth); Orange curve: *σ* =1 (orange; rugged). Dotted vertical line (corresponding to zero fitness difference) indicates critical value of correlation; above this critical value, switching between rugged landscape pairs (*σ* = 1) leads to larger fitness gains than evolution in a static landscape. B. Heatmap showing regions of parameter space (ruggedness *σ*, inter-landscape correlation) where switching leads to higher (black) or lower (white) fitness than evolution in a static landscape. C. Identical to panel A, but curves are shown for 5 (blue), 11 (red) and 501 (black) total evolutionary steps. *σ*=1 for all curves. D. Collateral fitness change, ranging from blue (less fit) to red (more fit), for populations adapted to alternating environments A and B as a function of ruggedness (*σ*) and inter-landscape correlation. Collateral fitness change is defined as the increase in average fitness in landscape B (relative to ancestor) associated with the steady state genotype distribution arising from adaptation to alternating A-B landscapes. *N* = 7 in all panels, but see also Figure S1. Error bars in panels A and C are ± standard error of the mean in the ensemble of landscapes.

## D. Fitness can be maximally increased in either static or alternating environments depending on the timescale

We find that adaptation to static environments typically occurs on a faster timescale than adaptation to alternating environments (Fig S3). As a result, the protocol yielding the highest average fitness may differ depending on the timescale over which the comparison is made. For example, on short timescales (5 total evolutionary steps; Fig 3C, blue), adaptation to static environments always leads to greater fitness gain, regardless of the correlation between landscapes. On moderate (11 total evolutionary steps; Fig 3C, red) to long (Fig 3C, black) timescales, however, we again see a range of positive correlations for which switching improves fitness–first only for highly correlated landscapes, and then eventually for a wider range of positively correlated landscapes. This result indicates that the optimal protocol for increasing fitness may depend on the chosen timescale; notably, recent results indicate that these timescales can also be tuned to maintain generalists successful in different environments (Sachdeva et al. 2020).

## E. Adaptation to alternating landscapes can lead to increased mean fitness even in anti-correlated landscapes

While we have so far focused on mean fitness defined in landscape A, either due to static 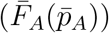 or alternating 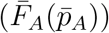 environments, we also asked how fitness in landscape B was modulated during adaptation. If adaptation occurs to a static landscape (A), the results are simple: the genotype adapted to A will on average exhibit increased (decreased) fitness in B when landscape B is positively (negatively) correlated with A. This scenario is reminiscent of collateral effects between different drugs, where increased resistance to one drug may be associated with either increased (cross resistance) or decreased (collateral sensitivity) resistance to a different (unseen) drug. In the case of alternating environments, however, the outcome is less clear *a priori*.

For smooth landscapes (*σ* = 0), we find that adaptation to the alternating landscapes leads to increased fitness in 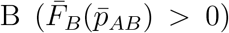 when the landscapes are positively correlated and decreased fitness when they are negatively correlated (Fig 3D). Nonzero epistasis shifts the boundary separating increased and decreased fitness toward negative correlations. As a result, switching leads to increased fitness in both landscapes for a wider range of correlations–even, counterintuitively, in cases where the landscapes are (weakly) anti-correlated. In the context of drug cycling, this result suggests that cross resistance is likely to arise following repeated cycling of two drugs, even when their fitness landscapes are anti-correlated (i.e. drugs induce mutual collateral sensitivity).

## F. Alternating between highly-correlated landscapes promotes escape from local fitness optima

To understand why switching between highly correlated landscapes can increase fitness relative to single landscape adaptation, we again represented adaptation on a simple (*N* = 4) network representing a particular pair of fitness landscapes (Fig S7). The choice of N=4 allows for a simpler visual interpretation of the results, and the relevant dynamics are qualitatively similar for a broad range of landscape sizes and sigma values (Fig S1, Fig S2). The landscape for environment A is characterized by multiple local maxima (Fig S7A, left panel), and in this example, the adaptation dynamics starting from the ancestral genotype are relatively simple, with only two paths possible (Fig S7A, right panel). With equal probability, the trajectory ends in one of two possible states, one of which is the global maximum.

If we now introduce rapid alternation with a second, positively correlated landscape (*ρ* = 0.8), the dynamics are much richer (Fig S7B). In this example, there is a single shared (local) maximum between the two landscapes (marked with red “+”), and adaptation to alternating environments eventually shepherds all trajectories to this shared maximum, which also happens to be the global maximum. As a result, alternating between landscapes leads to (on average) greater fitness increases than that achieved in static landscapes, where trajectories are split between local and global maxima. Intuitively, this example suggests that one advantage of rapid switching is that it dislodges trajectories from suboptimal local maxima–that is, switching between highly (but not perfectly) correlated landscapes provides a source of fluctuations that maximize the likelihood of finding globally optimal genotypes. This result is reminiscent of the observed “ratchet-like” mechanism of the *lac* operon in *Escherichia coli* (de Vos et al. 2015).

## G. Evolution in highly anti-correlated paired landscapes broadly samples genotype space resulting in reduced average fitness

We now return to dynamics in strongly anti-correlated landscapes, where shared maxima may be less likely to occur. To intuitively understand dynamics in this regime, we visualized the fitness landscape and evolutionary trajectories for a pair of simple (*N* = 4) anticorrelated landscapes (Fig S8). In this example, adaptation to the static landscape leads to considerably higher fitness than adaptation to alternating landscapes. Interestingly, we see that the genotype distribution remains broad, even for long times. In fact, the only genotypes that remain unoccupied (*p_i_* = 0) are those five that correspond to local minima in the A landscape. Including an additional step in landscape B leads to a similarly broad distribution, now with unoccupied genotypes corresponding to local minima of landscape B (Figure S5). In contrast to adaptation to single landscapes or alternating, positively correlated landscapes, the steady state distribution is not dominated by local fitness maxima but instead corresponds to broad genotype distribution and an associated decrease in average fitness.

## H. Adaptation to alternating landscapes is frequently dominated by presence or absence of shared fitness maxima

We hypothesized that the increased fitness in alternating landscapes is closely linked to the expected number of shared maxima between paired landscapes. To probe this hypothesis, we first estimated two quantities: 1) the fraction of local maxima that are shared between landscapes (specifically, the fraction of A-landscape maxima that also correspond to maxima in the paired landscape B) and 2) the fraction of landscape pairs that share at least one maxima. We estimated these quantities by simulating ensembles of landscapes and also developed semi-analytical approximations that reduce to simple evaluations of the cumulative distribution function (CDF) of a multivariate normal variable (SI). As intuition suggests, the fraction of shared maxima increases with correlation, both for smooth and rugged landscapes (Fig 4A). In addition, we estimated the fraction of landscape pairs in the entire ensemble that share at least one shared maximum (Fig 4B). Again we find that this quantity increases with correlation, but it does so much more rapidly for rugged landscapes. For smooth landscapes, the latter fraction increases gradually–and the curve is identical to that in (Fig 4A), a result of the fact that smooth landscapes have only a single (global) maximum.

**FIG. 4:**
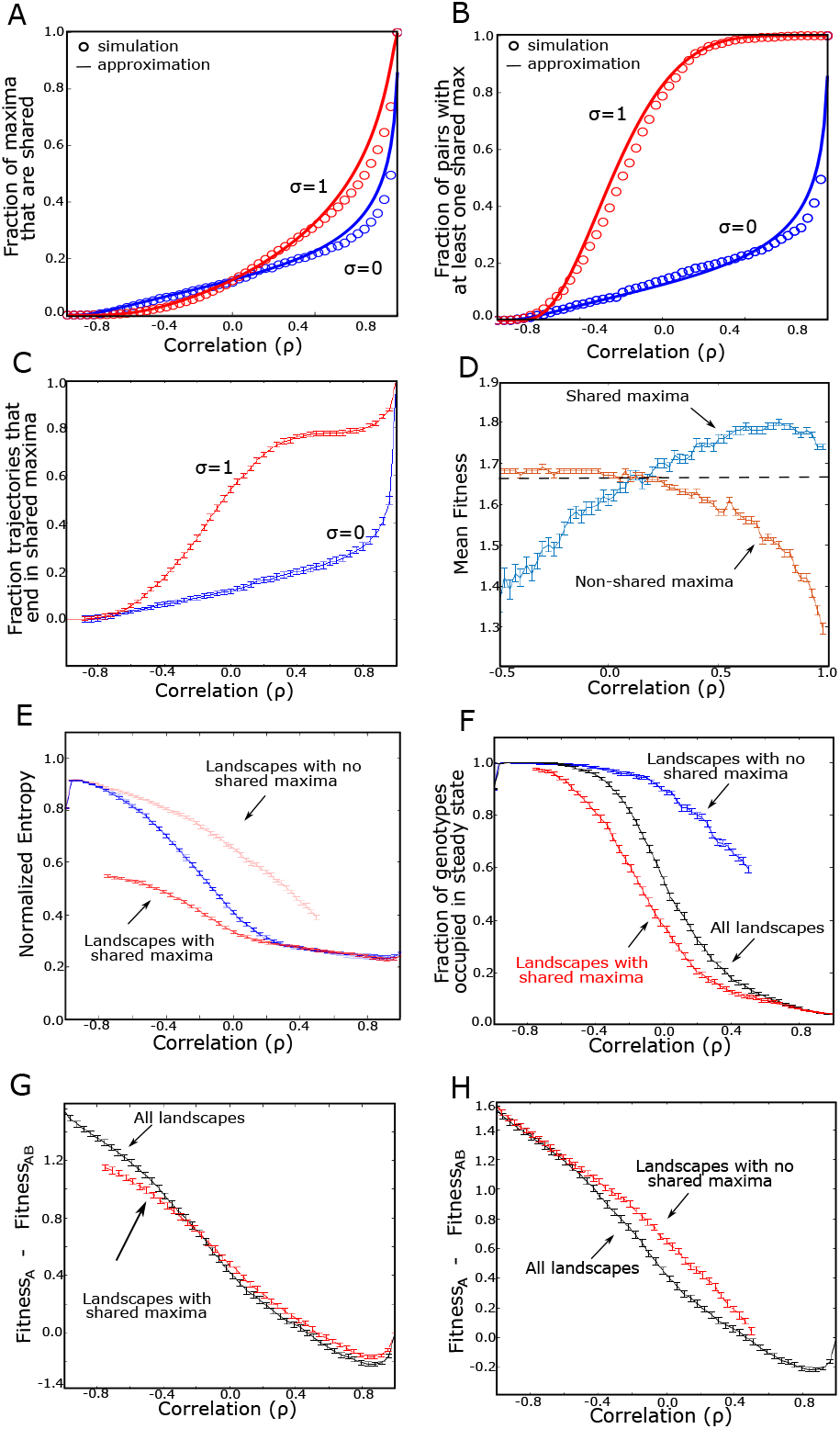
Evolution in alternating landscapes is frequently dominated by presence or absence of shared fitness maxima. A. Fraction of local maxima in landscape A that also correspond to a shared maxima in landscape B (*σ* = 0, blue; and *σ* = 1, red). B. Fraction of landscape pairs share at least one maximum. In panels A and B, circles corresponded to simulated landscapes and solid lines are semi-analytic approximations (see SI). C. Fraction of trajectories ending in a shared maximum as a function of correlation. D. Average fitness of shared maxima (blue) and average fitness of non-shared maxima (orange). Dashed line is average fitness of all local maxima in landscape A. E. Normalized entropy of the steady state genotype distribution following adaptation to alternating landscapes. Curves correspond to the full landscape pair ensemble (blue) and a reduced ensemble consisting only of landscapes that contain a shared maximum (red), bottom, and a reduced ensemble consisting only of landscapes with no shared maxima (red, top). The relative entropy is defined as *S*(*p*)/*S_max_* ≡ −(∑_*i*_ *p_i_* ln *p_i_*)/*S_max_*, where *p_i_* is the steady state probability of being in genotype *i* and *S_max_* is the entropy of a uniform distribution. F. Fraction of genotypes that have a nonzero probability of occupation in either the last A step or last B step at steady-state. Curves represent the paired landscape ensemble with no shared maxima (blue), the ensemble where every pair has at least one shared maximum (red), and the full ensemble (black). G. Difference in average fitness achieved in static and switching landscapes. Curves correspond to the full ensemble of paired landscapes (black) or a restricted ensemble that includes on those pairs that share a fitness maximum (red). H. Similar to panel F, with curves corresponding to the full ensemble (black) or a restricted ensemble that includes only those pairs with no shared fitness maxima (red). Error bars are ± standard error of the mean in the ensemble of landscapes. Error bars are ± standard error of the mean in the ensemble of landscapes. *N* = 7 for all curves, and *σ* = 1 for all curves in panels D-H.

To link these architectural properties of the landscapes with dynamics, we calculated adaptation trajectories under rapid switching of all paired landscapes in these ensembles (Fig 4C). For both smooth landscapes and negatively correlated rugged landscapes, the fraction of trajectories ending in a shared maximum closely mirrors the fraction of landscapes pairs that share a maximum. This correspondence suggests that under these conditions, when landscapes share a local maximum, the adapting system is likely to settle there. On the other hand, for positively correlated rugged landscapes, the likelihood of finding a shared maximum is relatively insensitive to correlation until *ρ* becomes quite large (> .80), when it rapidly increases (Fig 4C).

To further clarify the connection between fitness and shared maxima, we divided the local fitness maxima from landscape A into one of two categories: those that also correspond to a local maximum in landscape B, and those that do not. We found, somewhat counterintuitively, that the mean fitness differs for the two categories (Fig 4D). For negatively correlated landscape pairs, the fitness of shared maxima is less than that of non-shared maxima. By contrast, shared maxima in highly (positively) correlated landscapes have a higher mean fitness than non-shared maxima. In addition, there is a range of positive p where the fitness of shared maxima is also greater than the average fitness of maxima in a single A landscape (which corresponds to the *ρ* → 1 limit of the curve), offering an explanation for the fitness increase induced by alternating between highly correlated landscapes. Specifically, evolutionary trajectories typically settle into a single local maxima for adaptation to both static and positively correlated, alternating environments; however, for a range of highly (but not perfectly) correlated landscape pairs, the mean fitness of those shared maxima is greater than the mean fitness of local maxima in a single A landscape.

## I. Steady-state genotype distributions transition from narrow to broad as correlation is decreased

To further characterize steady state dynamics, we calculated the entropy of the steady state genotype distribution, defined as *S*(*p*)/*S_max_* ≡ −(∑_*i*_ *p_i_* ln *p_i_*)/*S_max_*, where *p_i_* is the steady state probability of being in genotype *i* and the expression is normalized by *S_max_* = *N* ln(2), the entropy of a uniform distribution (Fig 4E)–that is, a state where every genotype is equally probable. To capture dynamics associated with potential non-fixed point behavior, for this analysis we slightly modify the definition of steady state to be *p_i_* = (*p_A_ +p_B_*)/2, where *p_A_* is the steady state fitness following a step in landscape A (the previously used definition) and *p_B_* the fitness in the same steady state regime but following a step in landscape B (in words, we average over a full A-B cycle in the steady state). We find that as correlation (*ρ*) increases, the entropy of the system decreases, indicating that the dynamics are confined to an ever smaller set of genotypes–presumably those corresponding to shared maxima. Indeed, if we restrict the ensemble to only those landscape pairs that share a maximum, the entropy of the distribution is unchanged for highly correlated landscapes, suggesting that shared maxima dominate the steady state dynamics. By contrast, when landscape pairs are anticorrelated, restricting the ensemble to pairs *without* shared maxima closely approximates the results of the full ensemble, suggesting that dynamics in this regime are dominated by qualitatively different behavior. Consistent with changes in the entropy of the genotype distribution, we also find that correlation dramatically changes the fraction of genotype space occupied (with nonzero probability) in the steady state (Fig 4F). For highly correlated landscapes, only a small fraction of the total genotype space is occupied. By contrast, highly anti-correlated landscapes produce steady state distributions wherein all states are occupied with non-zero probability, suggesting ergodic-like behavior, consistent with the example in Fig S8. The fact that relative entropy remains less than 1 in this regime does indicate, however, that the distribution is not fully uniform.

Finally, in Fig 4G, we plot the difference in steady state fitness achieved in static vs alternating environments for both the full landscape pair ensemble (black) and for a reduced ensemble consisting only of landscapes with shared maxima (red). We find that the curves are nearly identical over a wide range of correlations *σ* > −0.4. Similarly, when correlation is strongly anticorrelated, fitness differences are similar between the full ensemble and the reduced ensemble with no shared maxima (Fig 4H). Taken together, these results provide evidence that adaptation in this model is frequently dominated by the presence or absence of shared fitness maxima, which in turn depends on the correlation between landscapes and landscape ruggedness.

## J. Clonal interference and slow switching reduce the impact of alternating between anticorrelated landscapes

Our idealized model neglects clonal interference, which could potentially impact the evolutionary dynamics (Gerrish and Lenski 1998). To investigate its potential impacts, we implement a phenomenological model previously used to estimate the effects of clonal interference (Tan and Gore 2012). Briefly, in the absence of clonal interference, the population can be treated as a *random walker* that steps to any nearby more fit genotype with equal probability. In order to simulate clonal interference, the population can be treated as a *greedy walker*, where the fixation probability of advantageous mutations is assumed to be proportional to s^*x*^, where *s* is the selective advantage and *x* is the phenomenological parameter. As x increases, the probability of stepping to more fit mutants continues to grow, simulating larger population sizes.

We find that small and moderate levels of clonal interference (*x* ~ 5) reduce the observed fitness differences between static and alternating protocols but lead to similar qualitative dynamics (Fig 5). However, as the population size gets large (*x* > 5) the fitness difference, genetic diversity and collateral effects due to switching become quite small; the impact of clonal interference is particularly large when landscape pairs are strongly anticorrelated.

**FIG. 5:**
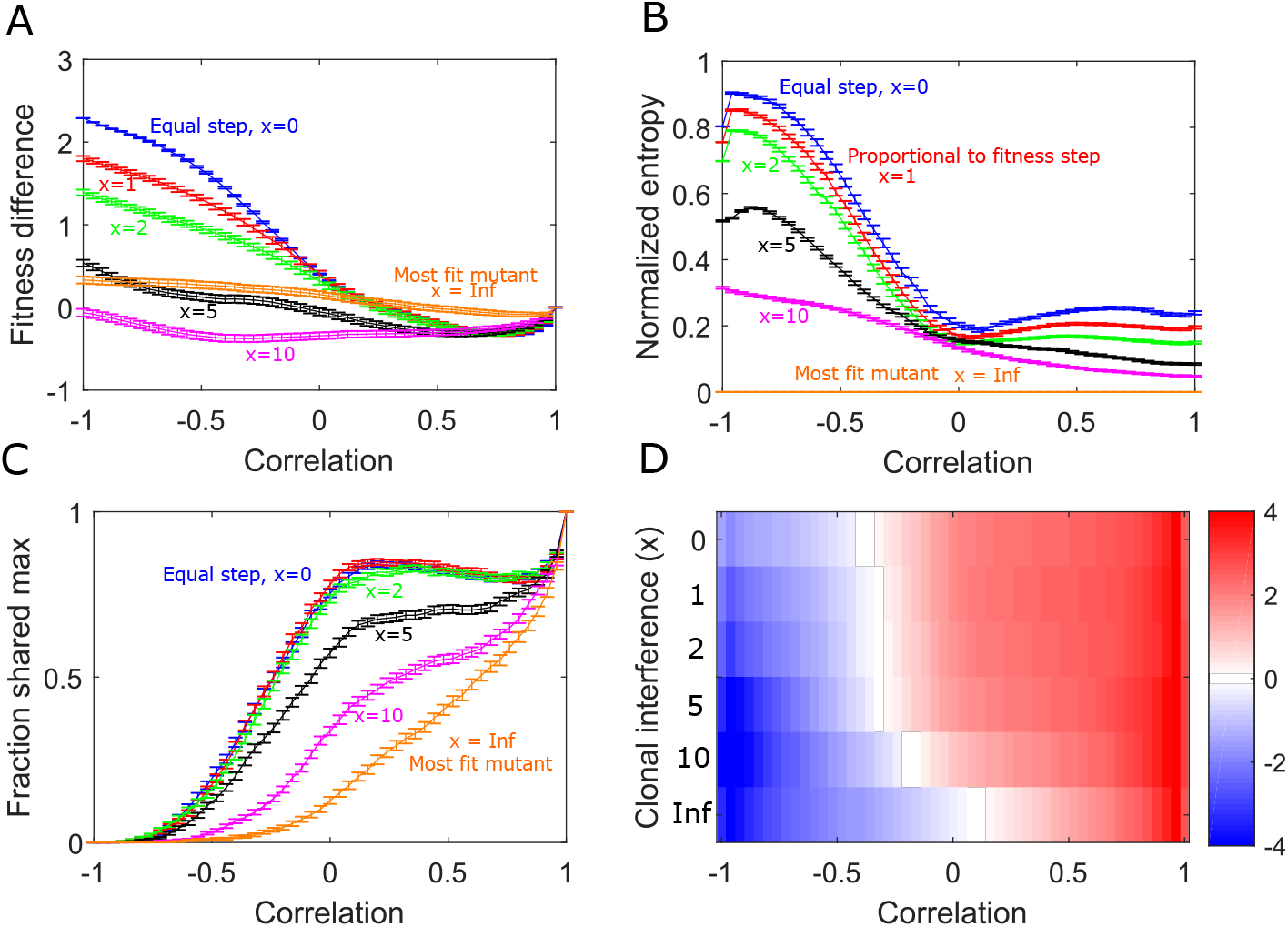
Clonal interference reduces the effects of alternating landscape evolution. A. Difference in average fitness achieved in static and switching landscapes. Curves correspond to different strengths of clonal interference (blue: random walker, *x* = 0, red: proportional walker, *x* = 1, green: *x* = 2, black: *x* = 5, magenta: *x* = 10, orange: *x infinite*, always steps to largest fitness neighbor). B. Normalized entropy of the steady state genotype distribution following adaption to alternating landscapes with different clonal interference. C. Fraction of trajectories ending in a shared maximum as a function of correlation with different clonal interference. D. Collateral fitness change, ranging from blue (less fit) to red (more fit), for populations adapted to alternating environments A and B as a function of clonal interference (*x*).

We next asked how the period of switching impacts the evolutionary dynamics. To do so, we varied the period of the switching (specifically, the number of consecutive steps taken in one landscape before switching) over approximately an order of magnitude (Fig 6). We find that small changes in the period–for example, doubling it from 1 step to 2– reduces the observed fitness differences and the normalized entropy, particularly for anticorrelated landscapes, but does not dramatically impact the likelihood of ending in a shared maximum or the collateral fitness changes (Fig 6).

**FIG. 6:**
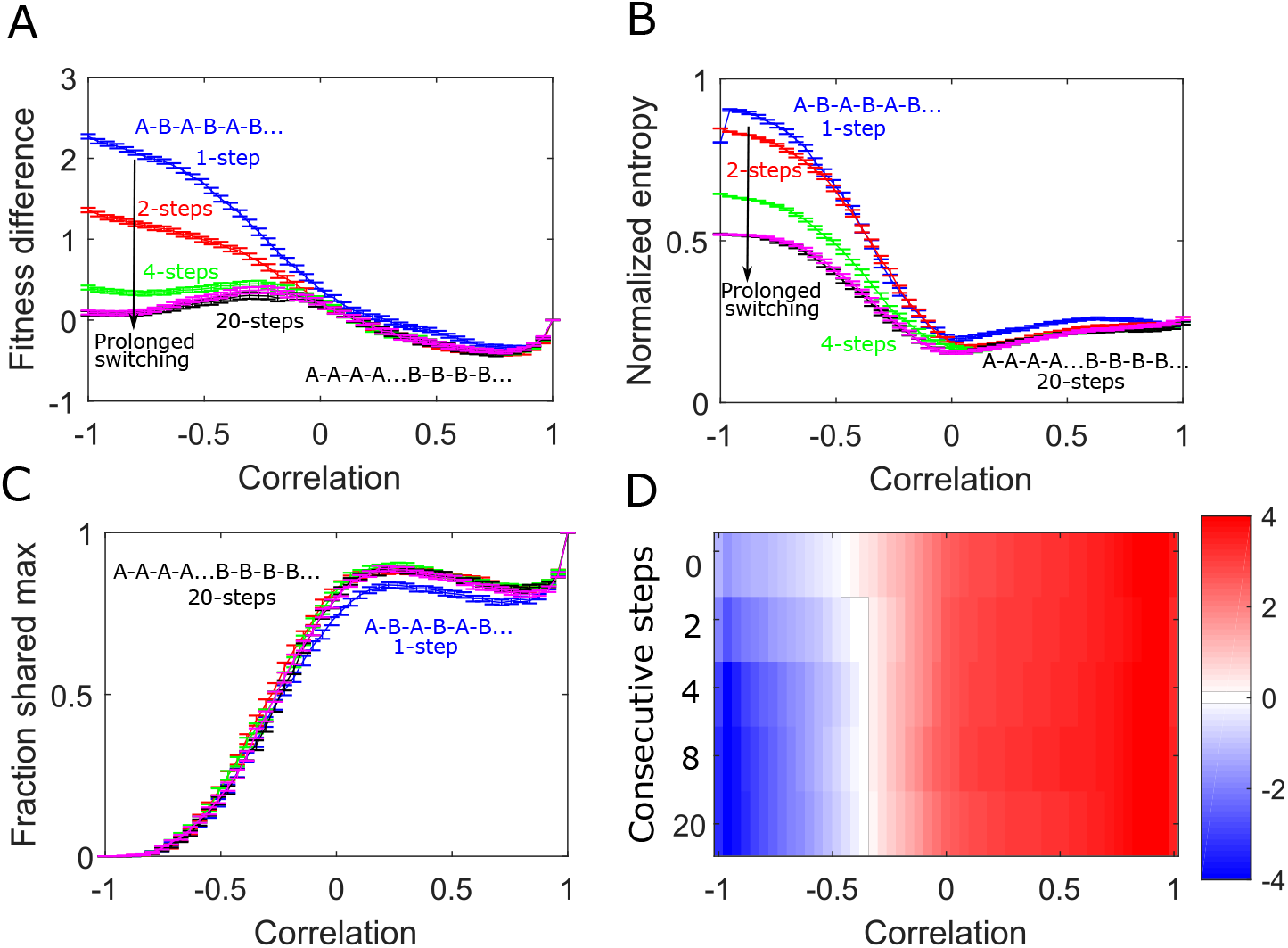
Consecutive steps in the same landscape before switching lessens the effects of alternating landscape evolution. A. Difference in average fitness achieved in static and switching landscapes. Curves correspond to different evolutionary steps taken in a landscape before switching (blue: 1 step, red: 2 steps, green: 4 steps, magenta: 8 steps, black: 20 steps). B. Normalized entropy of the steady state genotype distribution following adaption to alternating in landscapes with different switching periods. C. Fraction of trajectories ending in a shared maximum as a function of correlation with different switching periods. D. Collateral fitness change, ranging from blue (less fit) to red (more fit), for populations adapted to alternating environments A and B as a function of switching period.

## III. DISCUSSION

Our results indicate that both intra-landscape disorder (ruggedness) and inter-landscape fitness correlations impact fitness in rapidly alternating fitness landscapes. Compared with static adaptation, rapid switching can lead to increased or decreased fitness, depending on both the correlation between landscapes and level of intra-landscape ruggedness (i.e. epistasis). Perhaps most strikingly, switching between highly, but not perfectly, correlated rugged landscapes can increase fitness by promoting escape from local fitness maxima, increasing the likelihood of finding global fitness optima. Furthermore, rapid switching can also produce a genotype distribution whose fitness is, on average, larger than that of the ancestor population in both environments, even when the landscapes themselves are anti-correlated. Adaptation dynamics are often dominated by the presence or absence of shared maxima between landscapes. Rugged landscape pairs are increasingly likely to exhibit shared maxima as they become more positively correlated, and in turn, for landscapes with positive correlations, the mean fitness of these shared peaks is higher than that of non-shared peaks. By contrast, evolution in anti-correlated landscape pairs samples large regions of genotype space, exhibiting ergodic-like steady-state behavior that results in decreased average fitness. A simple phenomenological model suggests these results are robust to competition due to small and moderate clonal interference, however they disappear as population sizes grow excessively large. In addition, while prolonging the period of switching can alter the dynamics in anti-correlated landscape evolution, the fitness advantage conferred by alternating evolution in correlated landscape pairs is robust to the period of switching.

While our results are loosely inspired by antibiotic cycling, the model is highly idealized and certainly cannot make predictions that apply directly to clinical scenarios. At the same time, the simplicity and relative generality of the model means that it may be relevant for understanding the qualitative behavior of a wide range of systems, including evolution in antibodies (Burton et al. 2012), viruses (Rhee et al. 2010), and bacteria, where ratchet-like mechanisms for rapid adaptation have been observed experimentally (de Vos et al. 2015). Our model relies on the Strong Selection Weak Mutation (SSWM) limit and neglects potentially relevant dynamics that could arise due to horizontal gene transfer or population heterogeneity, which could potentially accelerate adaptation, particularly when switching between anticorrelated landscapes. While we also investigated an adapted model that accounts for clonal interference (Tan and Gore 2012), the model still assumes a homogeneous population, thus ignoring the genetic diversity necessary of clonal interference, and it neglects the possibility for deleterious or multiple simultaneous mutations to fix. In addition, we focus on small (typically *N* = 7) landscapes for tractability, and dynamics could differ for much larger landscapes.

It is important to note that we focus on paired landscapes characterized by a prescribed “global” correlation coefficient, but we do not investigate heterogeneity in the correlations at the single node level. In addition, the paired landscapes in our ensembles are constructed to share certain global features–like mean fitness–and are related by a prescribed interlandscape correlation, but they are not statistically identical. For example, the average number of local maxima can differ between landscape A and B, leading to different levels of evolved fitness for each landscape individually (Figure S6). This indicates that landscapes A and B have effectively different levels of epistasis, depending on the desired value of *ρ*, though these differences are most pronounced when A landscapes are very smooth (*σ* ≈ 0). These differences do not seem to be appreciably impacting fitness dynamics, as removing them by choosing a reduced ensemble (keeping only the B landscapes the exhibit similar fitness gains as A under static adaptation) does not appreciably modify the results (Figure S6). Nevertheless, it is possible that some of these results are specific to the exact manner in which correlated landscapes were produced; for example, in Figure 4D, the mean fitness for shared maxima equals that for unshared maxima at a small but nonzero value of *ρ* (rather than at *ρ* = 0), a counter intuitive result that may not hold when paired landscapes are generated by different algorithms. Indeed, it may be interesting to investigate switching dynamics using landscapes with different types of statistical similarities–for example, those that differ only in higher-order moments, or those that fully decouple landscape ruggedness and correlation (Wang and Dai 2019)). In fact, the results presented here are complementary to recent findings showing that environmental switching can enhance the basin of attraction for generalists, which are genotypes that are fit in multiple environments (Sachdeva et al. 2020; Wang and Dai 2019). While the focus of the work is different–and the timescale of environmental switching and the statistical relationships between landscape pairs differ in their model–our results similarly highlight the importance of shared landscape maxima in determining adaptation dynamics. Future work may aim to further elucidate the evolutionary impacts of varying timescale, ordering, and temporal correlations in landscape dynamics. In the long run, we hope results from idealized models like these offer increased conceptual clarity to complement the rapidly evolving experimental approaches for mapping landscape dynamics in living organisms.

## IV. METHODS

### A. Construction of the landscapes

We use the “rough Mt. Fiji” landscape model (Aita and Husimi 1998; Neidhart et al. 2014; Tan and Gore 2012) where each genotype is represented by a binary sequences of length N. The fitness of the ancestor genotype (0,0,0…0) is set to zero and the fitness *f_i_* associated with a single mutation at mutational site *i* is drawn from a uniform distribution on the interval [-1,1]. The fitness associated with multiple mutations is additive, and landscape ruggedness is incorporated by adding to the fitness of each genotype *j* a fixed, random variable *ξ_j_* drawn from a zero-mean normal distribution with variance *σ*^2^.

To create paired fitness landscapes, we represent each landscape A as a vector 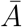 of length 2^*N*^, which we center and rescale to achieve a zero mean, unit variance vector. Then, we generate a Gaussian random vector 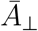 (also with zero mean and unit variance) and subtract from 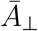 its projection onto 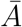, making 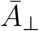 orthogonal to 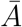; by construction, this vector corresponds to a landscape whose fitness values are, on average, uncorrelated with those of landscape A. It is then straightforward to generate a vector 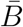, a linear combination of 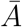 and 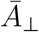, such that the fitness values of landscapes A and B are correlated to a tunable degree −1 ≤ *ρ* ≤ 1, where *ρ* is the Pearson correlation coefficient between the two vectors 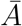 and 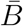. At the end of this procedure, we rescale 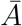 and 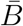 so that both have mean and variance equal to that of the original A landscape.

### B. Evolution on the landscapes

The SSWM assumption allows the evolutionary trajectories to be modeled as a Markov chain (Durrett and Durrett 1999; Nichol et al. 2015). We follow the “random move SSWM model”, which says that the probability of transitioning between adjacent genotypes *i → j* is given by *T_ij_* = 1/*m*, with *m* the total number of *i*-adjacent genotypes with fitnes greater than that of *i* in the given environment. Each environment (A or B) has its own transition matrix, which we designate as *T_A_* and *T_B_*. Evolution in environment A is then given by

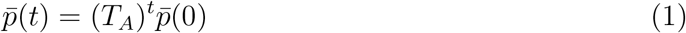

with 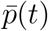 the vector whose *i*^th^ component is the probability to be in genotype *i* at time step *t*. We refer to the steady state (*t* → ∞) limit of this process as 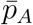. Similarly, we can describe rapidly alternating landscapes (A-B-A-B…) with

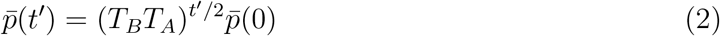

with *t*′ ≡ 2*t* an even time step. We refer to the steady state (*t* → ∞) limit of this process as 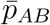. In practice, we define steady state using the condition 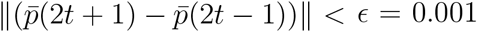. In words, we require the change in 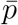 between consecutive steps in environment *A* to be sufficiently small. To facilitate comparison with static evolution in landscape A, we always end the process after a step in landscape A, meaning there are always an odd number of steps. Ending instead in landscape B results in qualitatively similar behavior, though the fitness is often shifted, indicating that a single step in A or B–even in steady state–can lead to significant changes in fitness S4.

## SUPPLEMENTAL MATERIAL

The Supplemental Material contains semi-analytic approximations for the number of local maxima in a single landscape and the probability of shared maxima in paired landscapes. It also includes seven supplemental figures (S1-S7).

### Semi-analytical approximations to describe local maxima

Dynamics in alternating environments are often impacted by the presence of shared local maxima. Here we derive semi-analytical approximate expressions for several key quantities. While exact expressions are difficult to obtain, even for the simple model used here, we derive below several approximations that involve cumulative distribution functions (CDFs) for common distributions (e.g. multivariate normal) and/or Gaussian-like integrals that can be easily calculated numerically.

### Number of local maxima in a single landscape

Rugged landscapes (*σ* > 0) potentially have multiple local maxima. To approximate the expected number of local maxima, consider a landscape with *N* loci so that each genotype has a total of *N* nearest neighbor genotypes. In a purely additive (*σ* = 0) landscape, a mutation in gene *i* changes the fitness by an amount *ϵ_i_* drawn from a uniform distribution [0,1]. In a rugged landscape, the fitness of each genotype also contains an additive contribution from epistasis–in this case, a zero-mean, normally distributed random variable with variance *σ*^2^. The fitness of each genotype is therefore a sum of (up to *N*) uniform variables and a single normally distributed variable.

To estimate the expected number of maxima in a landscape, consider a particular genotype with a fixed fitness *f = f_ϵ_ + f_σ_*, where *f_ϵ_* is the total fitness contribution from any mutations and *f_σ_* is the contribution from epistasis. The genotype will have *N* neighbors, each differing by a single mutation; the fitness of neighbor i has a fitness of the form *f_i_ = f_ϵ_ + ϵ_i_ + σ_i_*, where *ϵ_i_* is a uniform random variable accounting for adding or subtracting one mutation, and *σ_i_* is a normal random variable accounting for epistasis. The *f_ϵ_* term is a fixed value-the same as for the focal genotype. In a statistical ensemble of such neighbors, the probability that *f > f_i_*, that is, the probability that the genotype in question has a higher fitness than one particular neighbor is given by

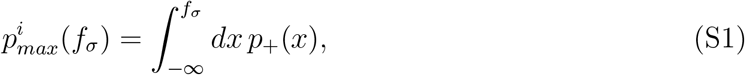

where *p*_+_(*x*) is the probability density function (pdf) for the sum *σ_i_ + ϵ_i_*. Since the pdf for a sum of random variables is given by the convolution of their individual pdfs, we have

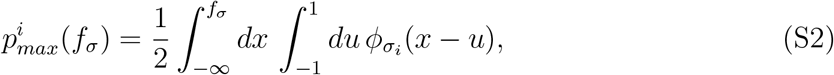

where *φ_σ_*(*x*) is the pdf of a zero-mean normal variable with variance *σ*^2^,

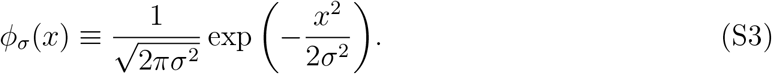

Equation S2 can also be written as

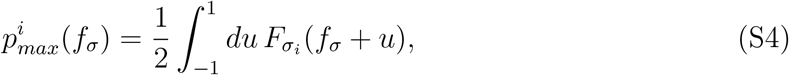

where *F_σ_i__*(*x*) is the cumulative distribution function for the variable *σ_i_* (and in this case, each *σ_i_* is a zero-mean normal variable with variance *σ*^2^). The integral above can be written as a linear sum of error functions, though it is somewhat cumbersome and we do not write it out here.

The probability that the genotype in question is a local max–that is, has a fitness larger than each of its *N* nearest neighbors-is approximately

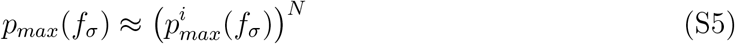

where we have assumed that each neighbor can be treated as independent from the others. The average probability that a genotype is a local maximum is then given by integrating over the distribution for *f_σ_*,

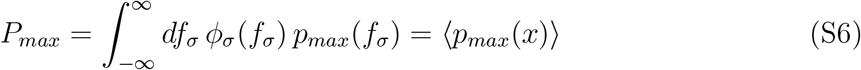

where brackets 〈·〉 represent an average over a normal distribution with variance *σ*^2^. If we assume that the 2^*N*^ different genotypes in a landscape are approximately independent, the expected number of maxima is then given by

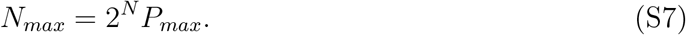

Equation S7 is difficult to evaluate analytically but easy to solve numerically, and the approximation closely matches results from randomly generated landscapes (Figure 3). For small epistasis (*σ* ≪ 1), we can expand *p_max_*(*f_σ_*) about the average (*f_σ_*) to arrive at the approximation

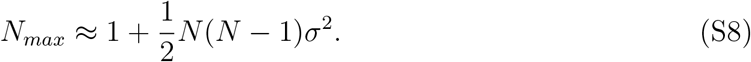

Similarly, for large epistasis (*σ* → ∞), we have *P_max_* ≈ (*N* + 1)^-1^; intuitively, all genotypes in a local neighborhood (the focal genotype and its *N* nearest neighbors) are equally likely to be the maximum, and the expected number of maxima therefore approaches *N_max_* = 2^*N*^/(*N* + 1).

### Shared maxima between correlated landscapes

Given that a particular genotype corresponds to a local maximum in landscape A, we would like to estimate the probability that it is also a maximum in the paired landscape B. To do so, consider a genotype that is a local maximum in landscape A. Let the fitness of that genotype be *a*_1_ and the fitness of its *N* nearest neighbors be *a*_2_ > *a*_3_… > *a_N_*_+1_, where we have labeled the neighbors according to their ranked fitness. We would like to calculate the conditional probability *p*(*b*_1_,*b*_2_..*b*_*N*+1_|*α*_1_,*α*_2_..*α*_*N*+1_) that describes the fitness values {*b_i_*} of the corresponding genotypes in landscape B, which is correlated with landscape A with correlation *ρ*.

In the limit of large epistasis (*σ* → ∞), the fitness variables {*a_i_*} and {*b_i_*} are jointly distributed normal variables with mean 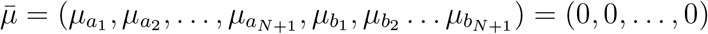 and covariance matrix

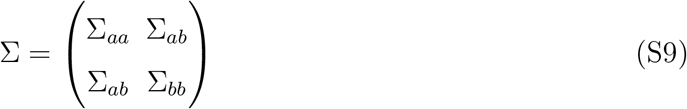

made of (*N* + 1) × (*N* + 1) sub-matrices

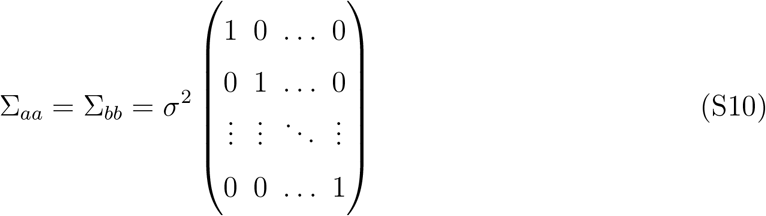

and

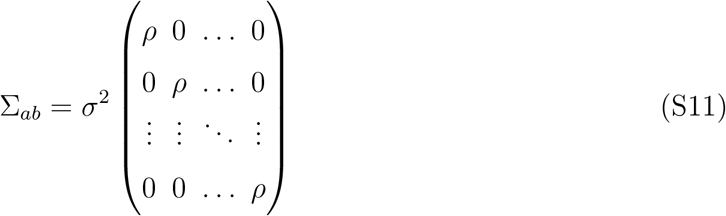

The matrix ∑_*aa*_ (∑_*bb*_) describes the covariance relationships between the focal genotype in landscape A (B) and each of its *N* nearest neighbors. The matrix ∑_*ab*_ describes the covariance between fitness values for the local neighborhood of *N* + 1 genotypes in landscapes A and B. If we treat the fitness values {*a_i_*} as fixed and the fitness values {*b_i_*} as random variables, the conditional probability *p*(*b*_1_, *b*_2_,… *b*_*N*+1_|*a*_1_, *a*_2_,… *a*_*N*+1_) is also normally distributed, with mean vector 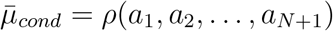 and covariance

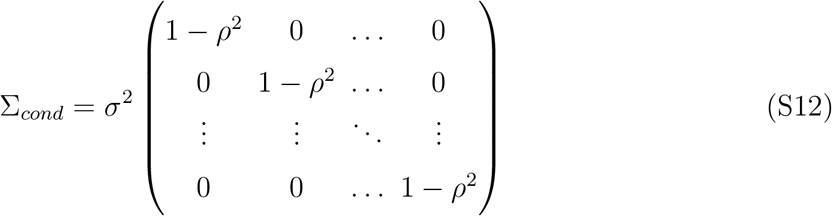

We would like to know the probability of *b*_1_ corresponding to a local maximum in landscape B given that *a*_1_ corresponds to a local maximum in landscape A. To do so, we consider the N variables *δ_i_ = b_i_ − b*_*i*−1_, whose distribution (conditioned on a specific set of values {*a_i_*}) is a multivariate normal with mean 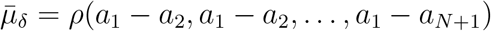 and covariance

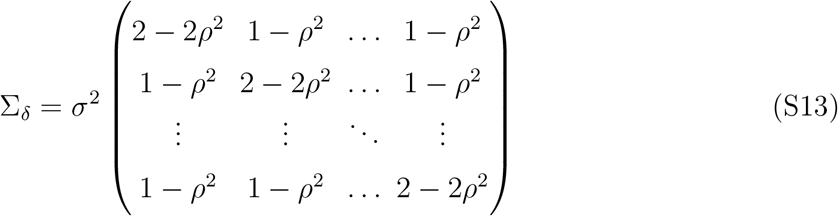

Hence, if we are given a specific set of fitness values {*a_i_*}, with *a*_1_ the maximum of the local fitness neighborhood in landscape A, the probability that the fitness is also a maximum in the B landscape is given by

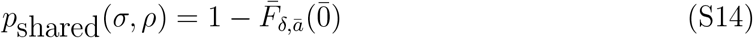

where 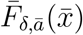 is the cumulative distribution function (CDF) for the multivariate normal with mean 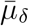 and covariance ∑_*δ*_ (conditioned on a set of values 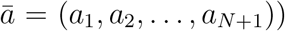) and 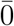 is the zero vector. Specifically, we have

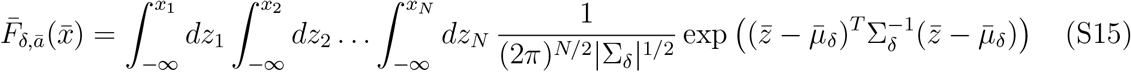

where *x_i_* is component *i* of 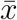. While there is no closed expression for the CDF of a multivariate Gaussian, there are many algorithms to rapidly calculate it numerically, and many scientific computing platforms even have built-in functions for this purpose.

To complete our approximation, we must choose specific values of the fixed variables {*a_i_*} on which the approximation is conditioned. In what follows, we consider two choices that lead to approximate expressions in the limits of of large and small epistasis.

In the limit of large epistasis, the fitness values in the local neighborhood {*a_i_*} are uncorrelated, Gaussian variables with variance *σ*^2^. We therefore choose *a_i_* to be the expected value of the *i*-th largest value in a sample of Gaussian variables (i.e. an order statistic (David and Nagaraja 2004)); without loss of generality, we assume the variables have mean zero. While there is no analytical expression for the expected value of the order statistics for normal variables, multiple approximations have been proposed. Here, we use the approximation in Royston (1982), which gives for our *N* +1 fitness variables

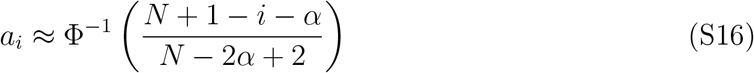

where Φ^−1^ is the inverse CDF for the unit Gaussian and *α* = 0.375 (we note that the order statistics can be calculated numerically to high precision, which slightly improves the approximation). Therefore, in the large *σ* limit, we have

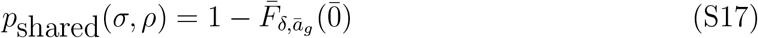

where the *i*-th component of 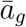 is given in Equation S16.

In the limit *σ* → 0, the fitness values in the local neighborhood {*a_i_*} are uncorrelated variables drawn from the uniform distribution [-1,1] (where again we choose a zero mean distribution without loss of generality). In this case, the expected value of the order statistics for uniform variables leads to

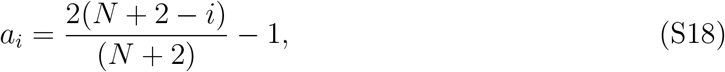

and our approximate expression is therefore

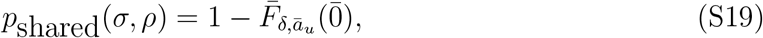

where the *i*-th component of 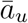 is given in Equation S18.

Finally, the fraction of landscape pairs that share at least one maximum is given by

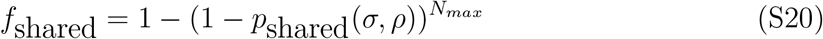

The approximations in Equations S17-S20 are not exact, but we find that they agree quite well with results from simulated landscapes in the *σ* = 0 (low epistasis) and *σ* = 1 (high epistasis) cases (Figure 4A and 4B).

---

**FIG. S1:**
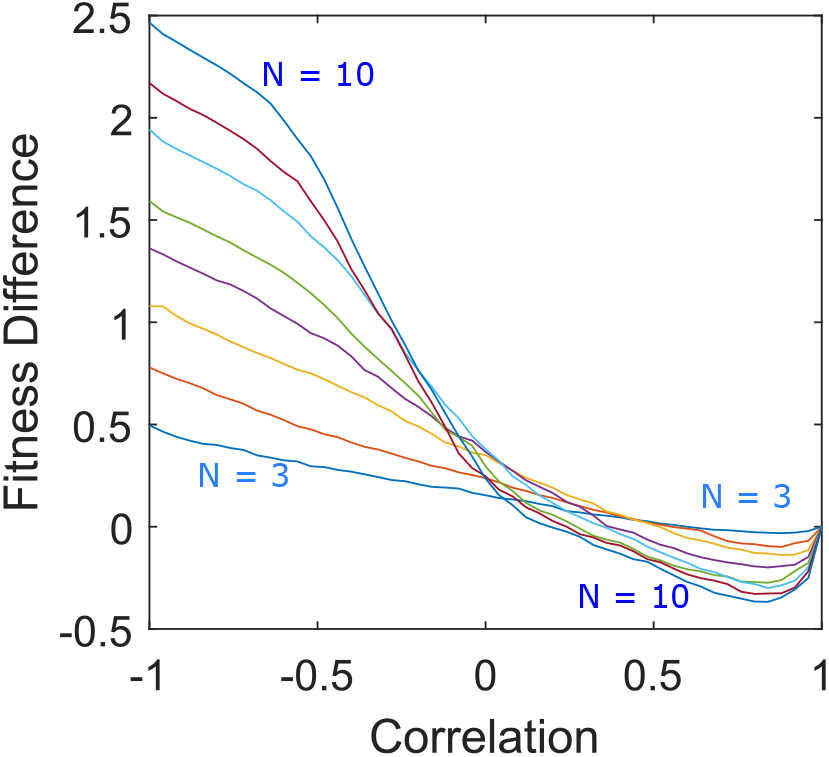
Rugged landscapes of different sizes show qualitatively similar changes in fitness as a function of correlation. Difference in average fitness (at steady state) between populations adapted to a single static landscape (landscape A) or rapidly alternating landscape pairs (A-B cycles) as a function of correlation between landscapes A and B. Average fitness is defined as the mean fitness of the steady state genotype distribution (which arises following adaptation to either static or switching protocols) measured in landscape A. Different curves range from *N* = 3 to *N* = 10, and *σ* = *N*/12 for each landscape to achieve relatively similar magnitudes of epistasis as *N* varies.

**FIG. S2:**
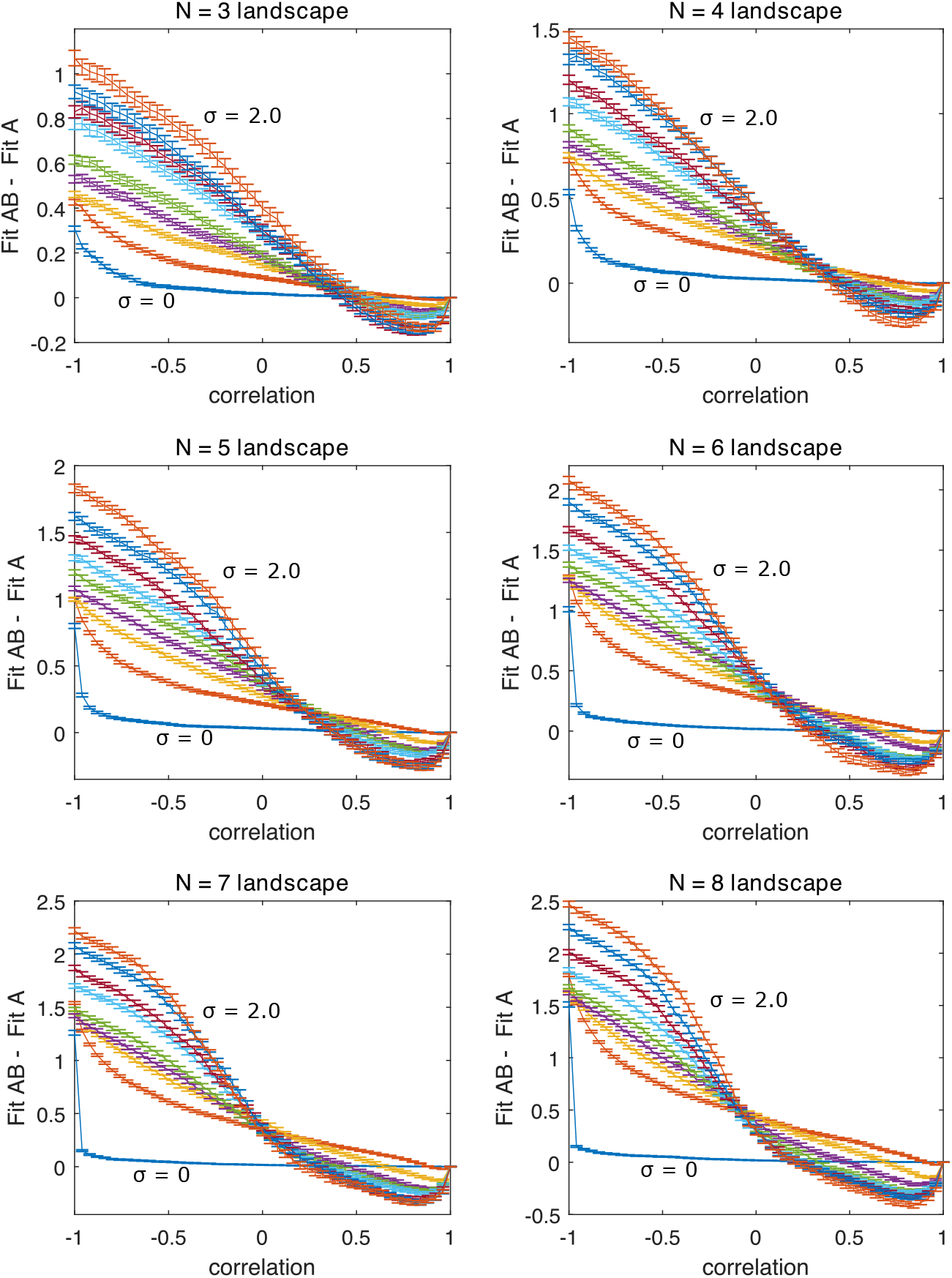
Landscapes of different sizes and sigmas show qualitatively similar results. Difference in average fitness (at steady state) between populations adapted to a single static landscape (landscape A) or rapidly alternating landscape pairs (A-B cycles) as a function of correlation between landscapes A and B. Average fitness is defined as the mean fitness of the steady state genotype distribution (which arises following adaptation to either static or switching protocols) measured in landscape A. Different curves range from *σ* = 0.0 (blue, labeled) to *σ* = 2.0 (orange, labeled) in increments of 0.25 for each landscape. Error bars represent the standard error of the mean for each simulation.

**FIG. S3:**
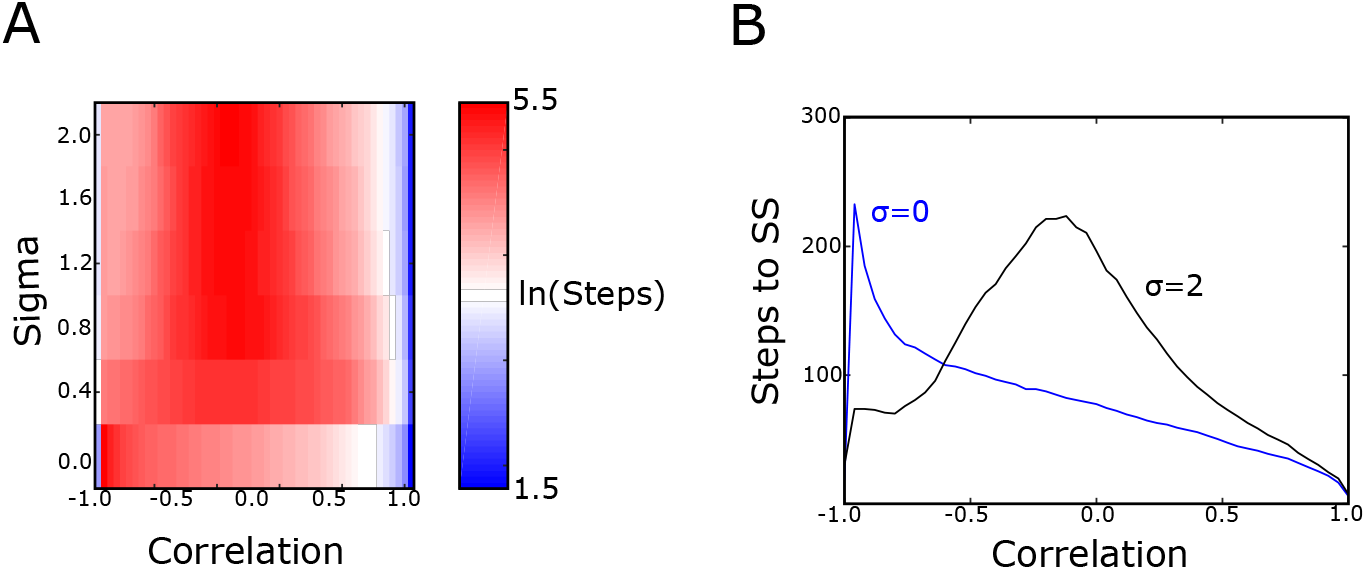
Adaptation to static and alternating environments approach steady state at different timescales. A. Number of time steps (log scale) until steady state for alternating landscapes of a given ruggedness (*σ*) and correlation (*ρ*). Full correlated landscapes (*ρ* = 1) correspond to static evolution in a single landscape. B. Example slices through panel A corresponding to *σ* = 0 and *σ* = 2.

**FIG. S4:**
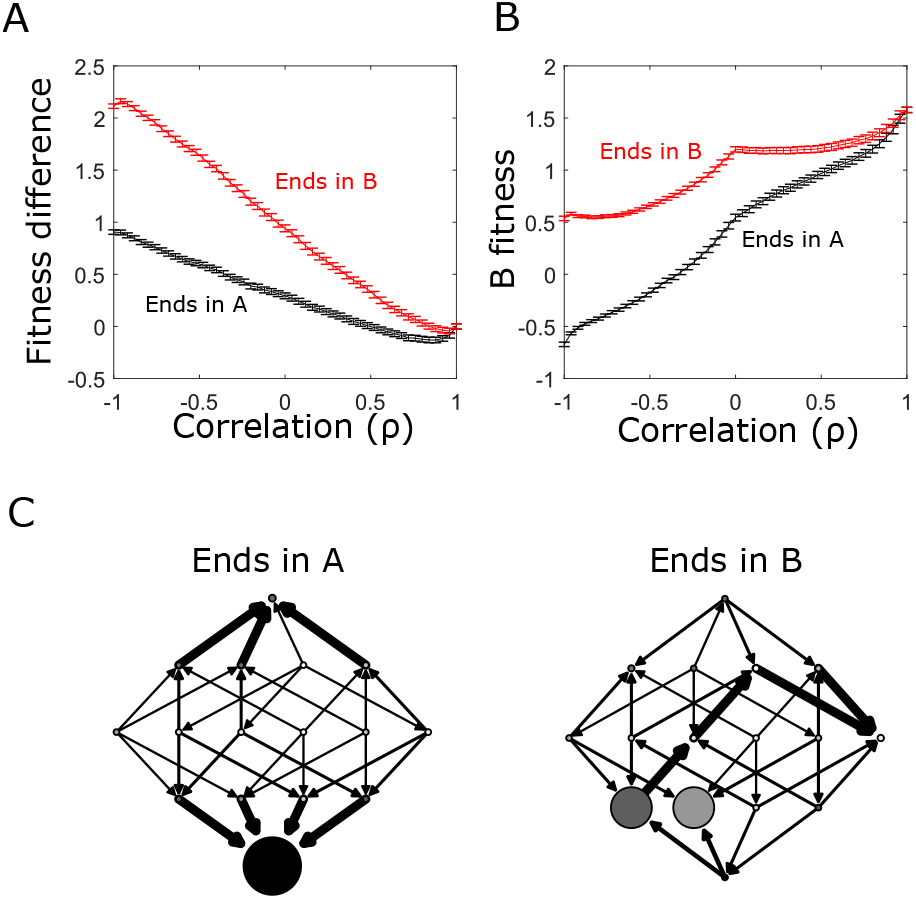
Adapted fitness depends on whether final step is taken in landscape A or B when landscapes are anticorrleated. A. Difference in average fitness (at steady state) between populations adapted to a single static landscape (landscape A) or rapidly alternating landscape pairs (A-B cycles) as a function of correlation between landscapes A and B. Average fitness is defined as the mean fitness of the steady state genotypte distribution (which arises following adaptation to either static or switching protocols) measured in landscape A. Curves correspond to steady state with a final step in landscape A (black) or a final step in landscape B (red). B. Collateral fitness change for populations adapted to alternating environments A and B as a function of inter-landscape correlation. Collateral fitness change is defined as the increase in average fitness in landscape B (relative to ancestor) associated with the steady state genotype distribution arising from adaptation to alternating A-B landscapes. C. Network representation of example fitness landscapes and transition probabilities following long-term adaptation to uncorrelated (*ρ* = 0) landscapes; adaptation ends either in landscape A (left) or B (right). *N* = 4 in all panels.

**FIG. S5:**
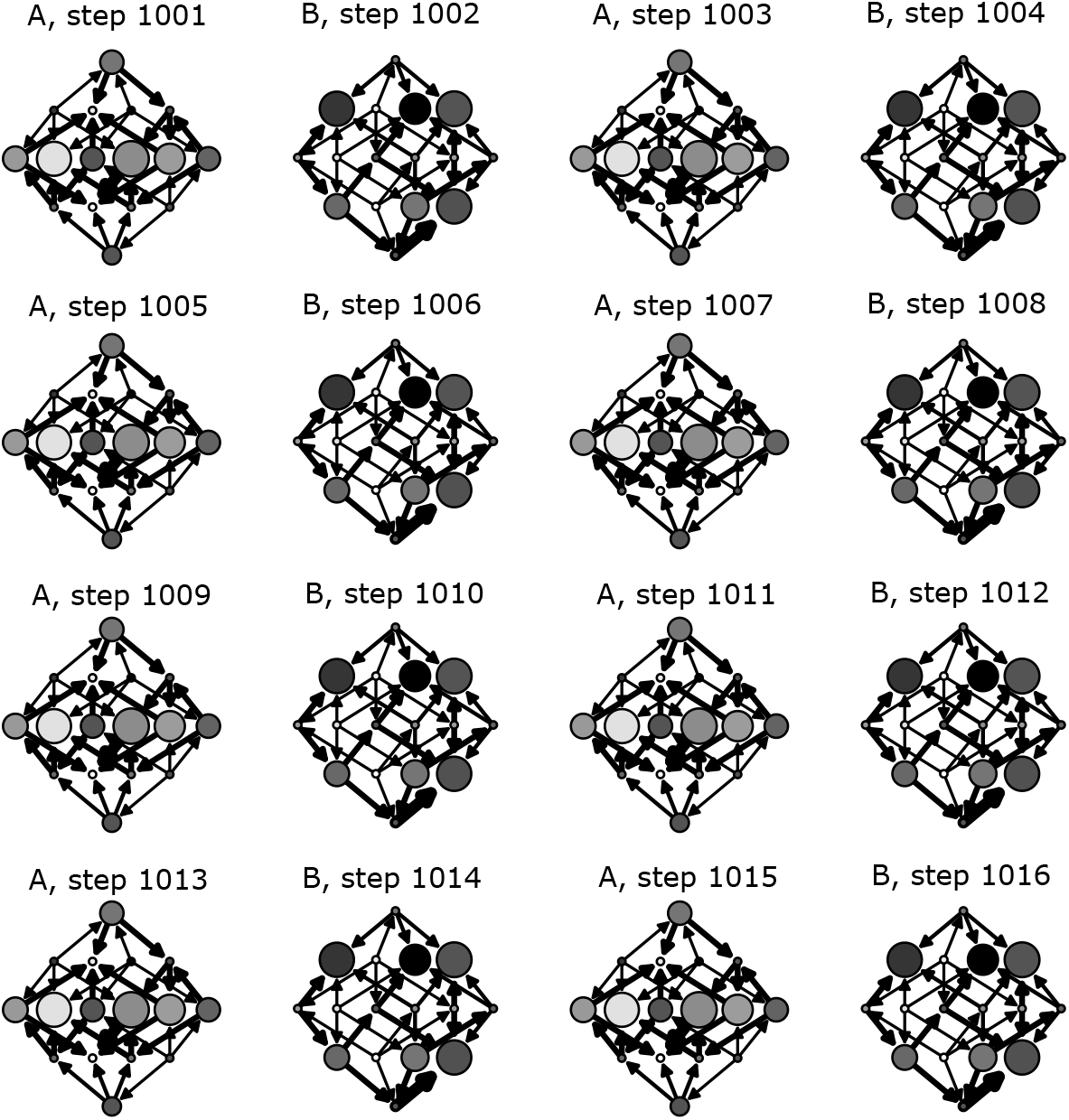
Adaptation to anti-correlated landscapes can produce cycles that sample large fractions of genotype space. Network representations of 16 consecutive steps in the steady state for paired landscape evolution with *ρ* = −0.88. Each circle represents a genotype (ancestral genotype at the top), with shading indicating the relative fitness of that genotype and size representing the occupation probability at that time step. Arrows represent transitions between genotypes that occur with nonzero probability and are accessible starting from the ancestor genotype. The width of the arrow represents the magnitude of the transition probability.

**FIG. S6:**
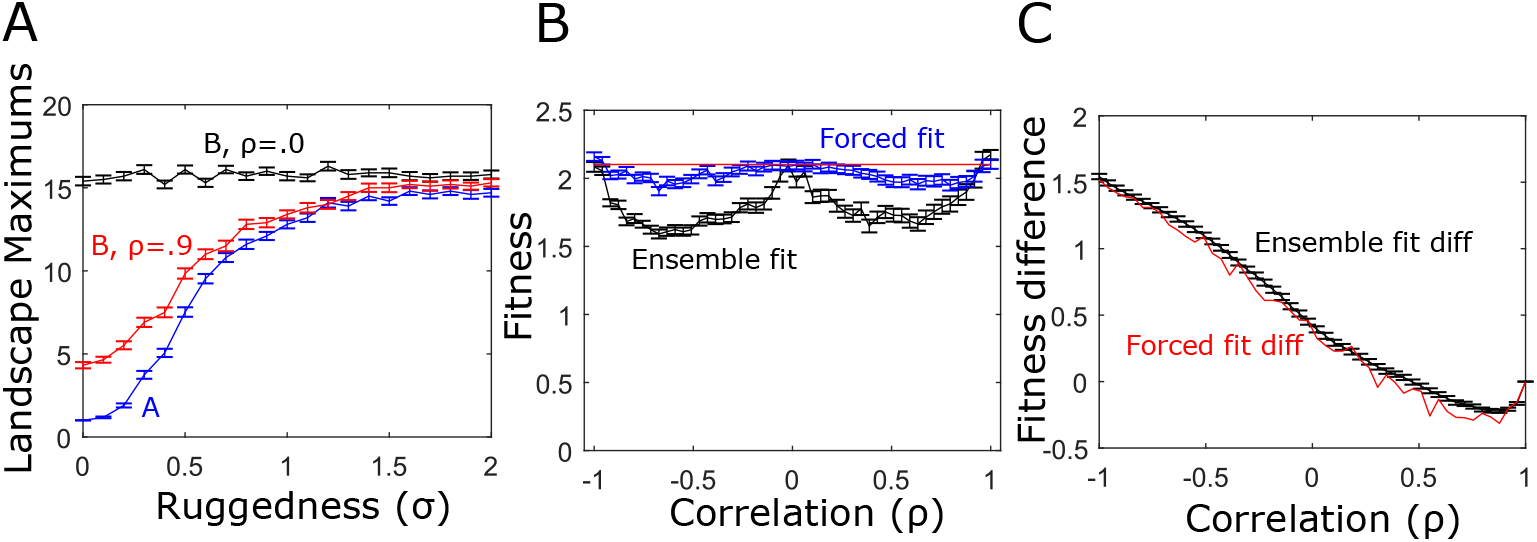
Statistical properties of landscape B differ from those of A but do not appreciably impact fitness differences between static and alternating landscapes. A. Average number of local maxima in landscape A (blue) and two different B landscapes correlated with A to different degrees (*ρ* = 0, black; *ρ* = 0.9, red). B. Evolved fitness following static adaptation to landscape A (red) or B (black). Blue curve is fitness in a reduced “forced fit” ensemble of B landscapes, which includes only those B landscapes that lead to similar levels of fitness as in landscape A. C. Fitness difference between static and switching environments for the full paired landscape ensemble (black) and for the reduced “forced fit” ensemble (black).

**FIG. S7:**
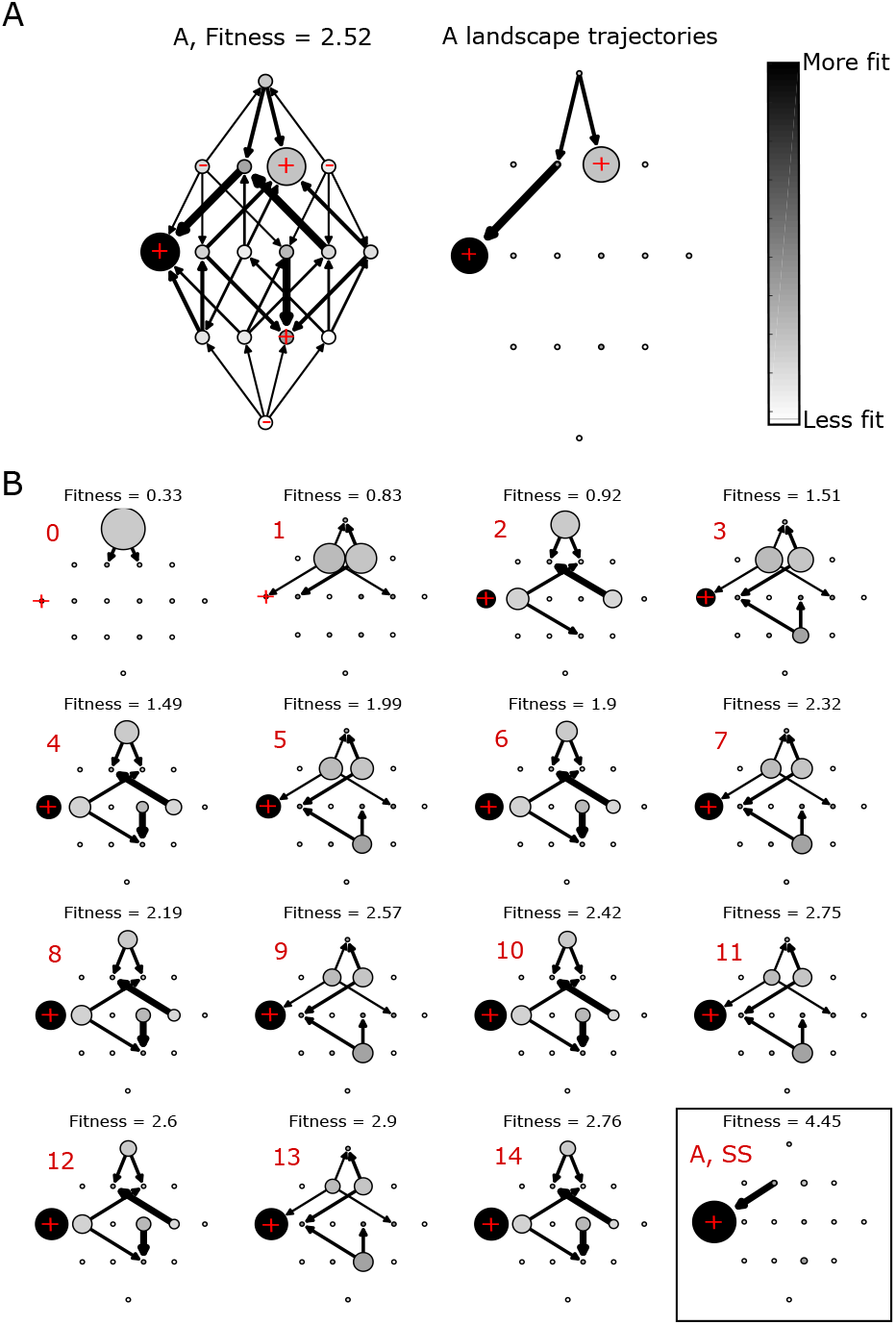
Evolutionary dynamics in alternating landscapes with positively correlated fitness peaks. A. Left panel: network representation of adaptation on a static landscape (environment A) of size *N* = 4. Each circle represents a genotype (ancestral genotype at the top), with shading indicating the relative fitness of that genotype and size representing the occupation probability in the steady state. Red + symbols mark genotypes corresponding to local fitness maxima. Arrows represent transitions between genotypes that occur with nonzero probability–that is, the entries of the transition matrix. The width of the arrow represents the magnitude of the transition probability. Right panel: same as left panel, but showing only transitions that occur during adaptation starting from the ancestral genotype (top circle). B. Network representations of adaptation (at different time points) in alternating landscapes with positively correlated fitness peaks. Red number above each landscape represents the current evolutionary time point (ranging from 0 to SS, indicating steady state of approximately 200 steps). Directed arrows represent possible transitions between genotypes based on the current genotype distribution (indicated by the circle sizes) and the current landscape (A or B). Average fitness at each time point (calculated over the current genotype distribution) are listed above each plot. Even numbered steps correspond to landscape A, odd to landscape B.

**FIG. S8:**
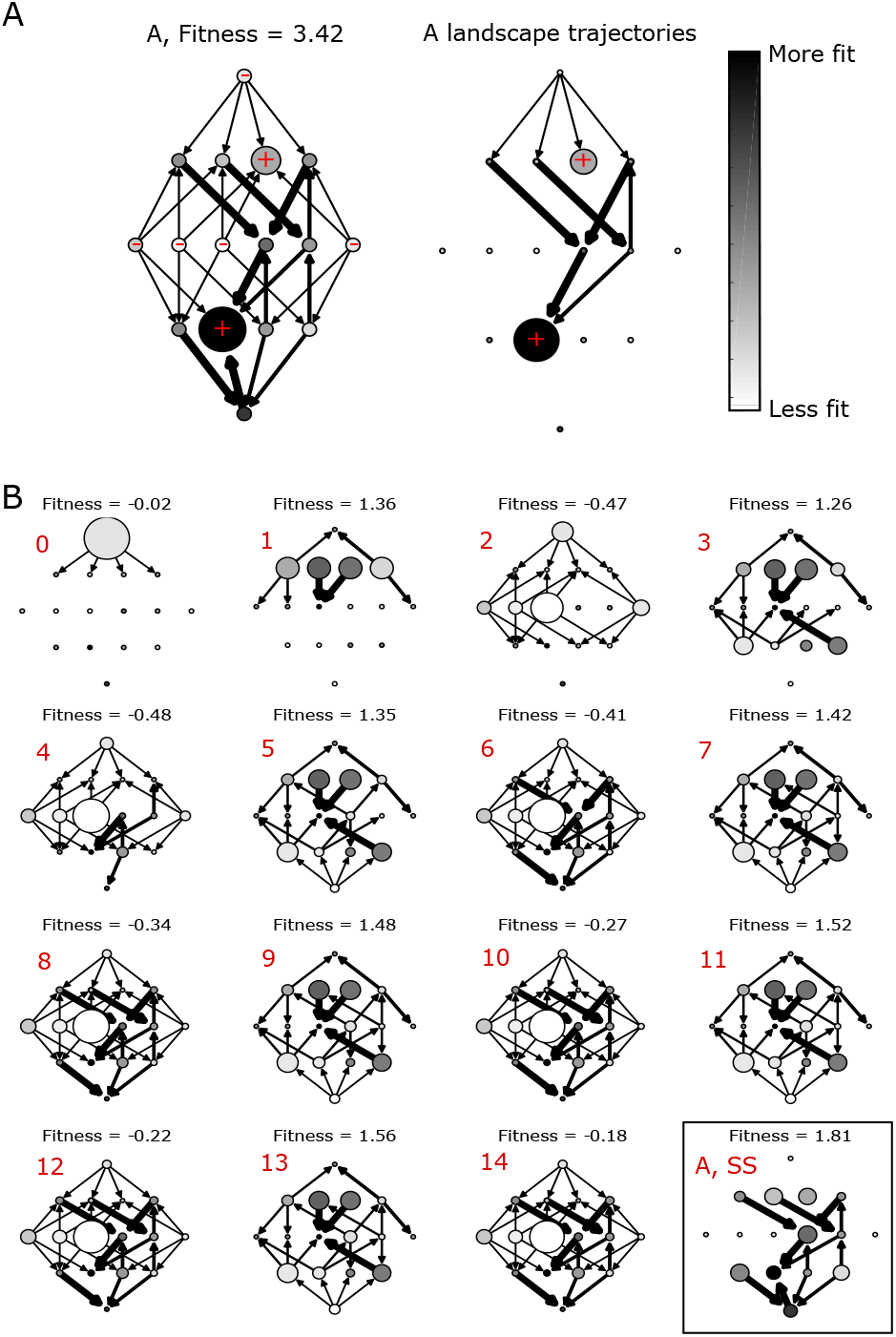
Evolutionary dynamics in alternating landscapes with negatively correlated fitness peaks. A. Left panel: network representation of adaptation on a static landscape (environment A) of size *N* = 4. Each circle represents a genotype (ancestral genotype at the top), with shading indicating the relative fitness of that genotype and size representing the occupation probability in the steady state. Red + symbols mark genotypes corresponding to local fitness maxima. Arrows represent transitions between genotypes that occur with nonzero probability. The width of the arrow represents the magnitude of the transition probability. Right panel: same as left panel, but showing only transitions that occur during adaptation starting from the ancestral genotype (top circle). B. Network representations of adaptation (at different time points) in alternating landscapes with negatively correlated fitness peaks. Red number above each landscape represents the current evolutionary time point (ranging from 0 to SS, indicating steady state of approximately 200 steps). Directed arrows represent possible transitions between genotypes based on the current genotype distribution (indicated by the circle sizes) and the current landscape (A or B). Average fitness at each time point (calculated over the current genotype distribution) are listed above each plot. Even numbered steps correspond to landscape A, odd to landscape B.

